# The Conformational Space of the SARS-CoV-2 Main Protease Active Site Loops is Determined by Ligand Binding and Interprotomer Allostery

**DOI:** 10.1101/2024.09.09.612101

**Authors:** Ethan Lee, Sarah Rauscher

**Affiliations:** Department of Chemical and Physical Sciences, University of Toronto Mississauga, Mississauga, ON, Canada L5L 1C6; Department of Chemistry, University of Toronto, Toronto, ON, Canada M5S 3H8; Department of Physics, University of Toronto, Toronto, ON, Canada M5S 1A7

## Abstract

The main protease (M^pro^) of SARS-CoV-2 is essential for viral replication and is, therefore, an important drug target. Here, we investigate two flexible loops in M^pro^ that play a role in catalysis. Using all-atom molecular dynamics simulations, we analyze the structural ensemble of M^pro^ in an apo state and substrate-bound state. We find that the flexible loops can adopt open, intermediate (partly open) and closed conformations in solution, which differs from the partially closed state observed in crystal structures of M^pro^. When the loops are in closed or intermediate states, the catalytic residues are more likely to be in close proximity, which is crucial for catalysis. Additionally, we find that substrate binding to one protomer of the homodimer increases the frequency of intermediate states in the bound protomer, while also affecting the structural propensity of the apo protomer’s flexible loops. Using dynamic network analysis, we identify multiple allosteric pathways connecting the two active sites of the homodimer. Common to these pathways is an allosteric hotspot involving the N-terminus, a critical region that comprises part of the binding pocket. Taken together, the results of our simulation study provide detailed insight into the relationships between the flexible loops and substrate binding in a prime drug target for COVID-19.

## 1 Introduction

Severe acute respiratory syndrome coronavirus 2 (SARS-CoV-2) is a betacoronavirus from the *Coronaviridae* family, of which SARS-CoV and Middle East respiratory syndrome coronavirus (MERS-CoV) are also members.^1^ SARS-CoV-2 spreads through human-to-human transmission and is responsible for the COVID-19 pandemic. During viral replication, two-thirds of the genome is translated into two polyproteins containing 16 nonstructural proteins (nsps).^2^ These polyproteins are cleaved by the papain-like protease and the main protease (M^pro^). M^pro^ cleaves the polyproteins at 11 sites, including two sites liberating itself. ^3,4^ Dimerization of M^pro^ is essential for its catalytic activity,^5,6^ as the N-terminal region of one monomer forms part of the substrate binding site of the other monomer. ^7^ Each M^pro^ monomer is 306 residues in length and comprises three domains. ^7,8^ Domains I and II are six-stranded antiparallel β-barrels that enclose the active site between them, while Domain III contains five helices and plays a role in dimerization. ^7,8^ The catalytic Cys-His dyad located in the active site of M^pro^ cleaves the polyprotein substrate. The first step in the reaction mechanism involves His_41_ accepting a proton from Cys_145_, which results in a charged thiolate and is followed by a nucleophilic attack on the carbonyl carbon atom of the peptide bond in the substrate by the S*γ* atom of Cys_145_.^9^

M^pro^ is a prime target for inhibition due to its importance in viral replication. ^7,10^ There are no human proteases that target similar cleavage sequences to M^pro^, which suggests that inhibitors would have a lower potential for toxicity. ^7^ To date, over 100 inhibitors of M^pro^ have been identified,^10^ and the Protein Data Bank (PDB) contains over 1200 structures of M^pro^. Furthermore, the FDA has approved a ritonavir-boosted nirmatrelvir protease inhibitor known as Paxlovid.^11^ Continued investigation of M^pro^ is crucial not only for enhancing the effectiveness and specificity of antiviral drugs, but also because several SARS-CoV-2 variants of concern have mutations in M^pro^,^12^ and mutations in M^pro^ can confer resistance to nirmatrelvir.^13^ Furthermore, a detailed understanding of the structure and function of M^pro^ will strengthen global preparedness against potential future pandemics where related viral proteases may be targets.

Most inhibitors of M^pro^ bind in or near its active site,^10^ which is surrounded by two mobile loops.^14,15^ Flexibility of the active site and the surrounding loops is functionally important because it allows for the accommodation of diverse substrates.^4^ Other viral proteases and enzymes have mobile active site loops that play critical roles in their function and dynamics.^16^ For instance, a study by Jupin et al. reported the switch action of an active site loop in an ovarian tumour-like protease/deubiquitinase. This loop samples either an open, intermediate, or closed state. These loop states are related to the protein’s function: the state of the loop determines whether the protein acts as a protease or a deubiquitinase. ^17^ Similarly, studies on HIV-1 protease showed two symmetric active site loops that populate open, intermediate, and closed states depending on the occupancy of the active site, as revealed by nuclear magnetic resonance (NMR) experiments and molecular dynamics (MD) simulations.^18,19^

MD simulations are a powerful tool for studying viral proteases, as well as mobile loops in enzymes more generally, providing complementary information to crystallographic and biophysical studies.^16^ Simulation studies of M^pro^ have advanced our understanding of the structure and dynamics of this enzyme in order to provide useful information for the development of viable inhibitors.^15,20–32^ The simulation study carried out in this work builds on the growing body of research into the functional motions of M^pro^. Specifically, we aim to describe specific conformational states of the active site in both protomers of the homodimer and quantify the extent to which information is transmitted between the active sites through interprotomer allostery.

Here, we focus on M^pro^’s flexible active site loops, presenting a combined 92.5 µs of all-atom simulation data on M^pro^ in different conditions and from multiple coronaviruses. With this extensive simulation data set, we demonstrate that the active site loops sample open, closed, and intermediate states, and the frequency of these states depends on the occupancy of the active site. We find evidence for interprotomer allostery: the frequency of the loop states in one protomer depends on the bound state of the opposite protomer. To further explore this allostery, we used dynamic network analysis to identify residues involved in interprotomer communication. Additionally, we find that the isolated monomer has a higher frequency of open states compared to the dimer, which may explain its weaker ligand binding.^6,26^ Finally, we compare M^pro^ of SARS-CoV-2, SARS-CoV, and MERS-CoV and find similarities between the three proteases in the dynamics of the flexible loops and the allosteric paths connecting the two active sites.

## 2 Results and Discussion

### 2.1 Assessment of Sampling, Equilibration, and Accuracy

In this study, we conducted MD simulations of the M^pro^ dimer with either one, two, or zero (apo) substrates bound. The complete list of all simulations performed is provided in Table 1. We define the bound systems that were simulated as follows: an M^pro^ dimer with two substrates bound (one per active site) is referred to as the *two-substrate system*. A complex with one substrate bound (where the dimer has one bound and one apo protomer) is referred to as the *one-substrate system*. Additionally, simulations were performed on M^pro^ as a monomer, with and without substrate, as well as on apo SARS-CoV and MERS-CoV M^pro^ dimers. To mimic the polyprotein cleavage site between nsp4 and the N-terminus of M^pro^, we used the same pentapeptide substrate as Suárez and Díaz. ^23^ This substrate contains a –P4–P3–P2–P1–P1’-sequence of -Ala–Val–Leu–Gln–Ser–, where cleavage would occur at the P1–P1’ site (Figure 1B and C, see SI Section 1.1 for more details).^23^ Equilibrated sections of the trajectories were determined using four metrics: root-mean-square deviation (RMSD), radius of gyration (R_gyr_), number of hydrogen bonds, and solvent accessible surface area (SASA). Convergence of these four metrics occurred at ∼0.5 µs (Figure 1D). Only the data after t = 0.5 µs were included in further analysis (see SI Section 1.2).

**Table 1:** Summary of M^pro^ MD simulations. The setup of substrate-bound systems is described in SI Section 1.1.

| Virus | Oligomeric State | Trajectory Length ($\mu\text{s}$) | Replicates | Bound State |
| --- | --- | --- | --- | --- |
| SARS-CoV-2 | Dimer | 1.5 | 5 | Apo |
| SARS-CoV-2 | Dimer | 4.0 | 5 | 2 Substrates |
| SARS-CoV-2 | Dimer | 4.0 | 5 | 1 Substrate |
| SARS-CoV-2 | Monomer | 3.0 | 5 | Apo |
| SARS-CoV-2 | Monomer | 3.0 | 5 | Substrate |
| SARS-CoV | Dimer | 1.5 | 5 | Apo |
| MERS-CoV | Dimer | 1.5 | 5 | Apo |

**Figure 1:**
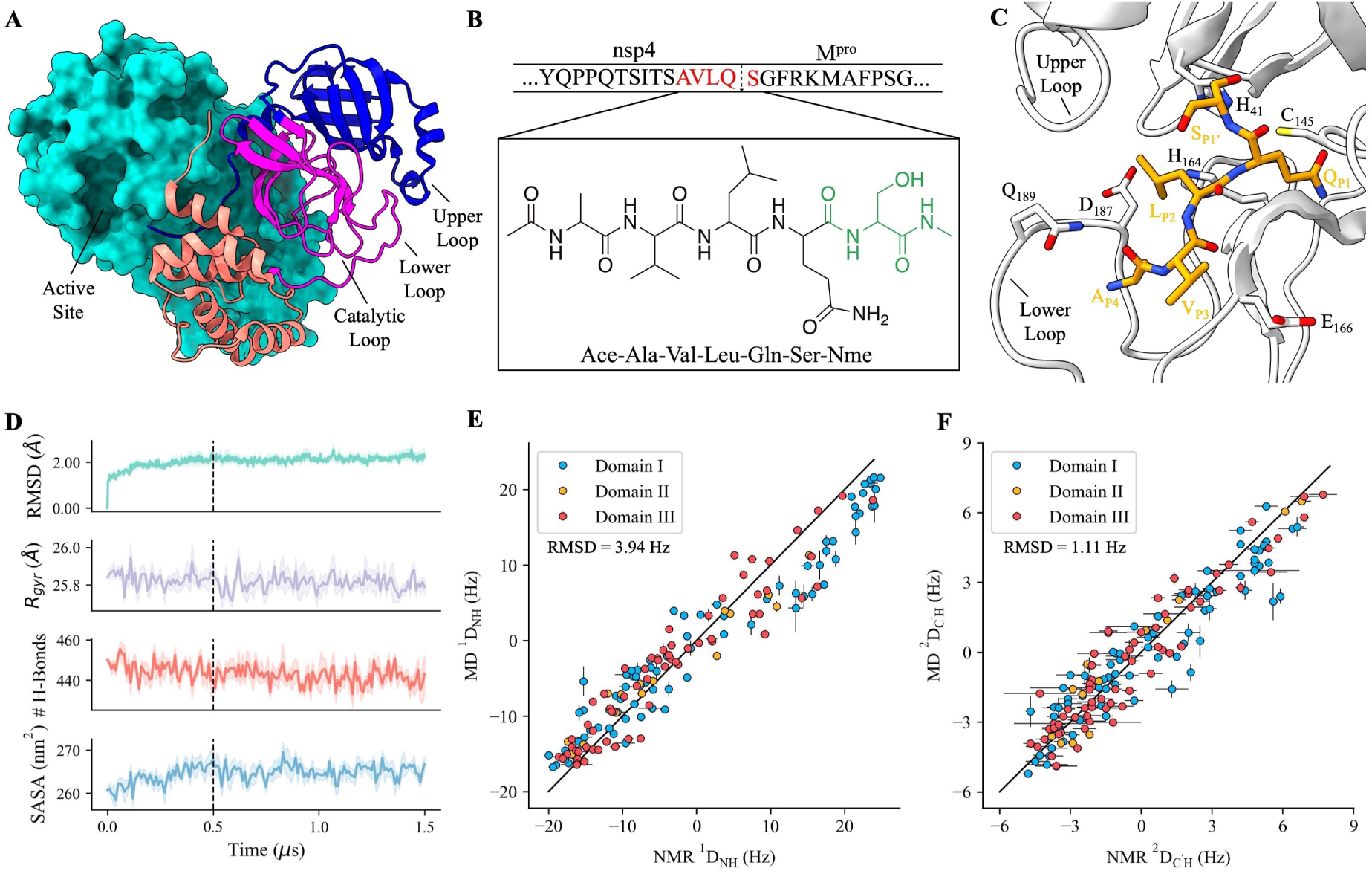
Simulation setup and validation. (A) Surface area and ribbon models show two M^pro^ protomers (PDB ID: 6XHU)^33^ The ribbon model is split into three domains: Domain I (blue, residues 8-101), Domain II (magenta, 102-184), and Domain III (orange, 201-303). The linker connecting Domains II and III (185-200) is also coloured magenta. We define three loops: the upper loop (residues 44-54), lower loop (184-194), and a loop in the active site that we refer to as the catalytic loop (162-174). (B) Pentapeptide substrate mimetic used in all simulations of the bound systems. The upper row shows the –P4–P3–P2–P1–P1’ sequence of -Ala-Val-Leu-Gln-Serin red, where cleavage occurs at the P1–P1’ site (arrow) between the sequence of nsp4 and M^pro^.^23^ The black portion of the peptide was taken from the N3 inhibitor in PDB 6LU7,^8^ while the green portion was constructed along the inhibitor backbone by Suárez and Díaz. ^23^ (C) The substrate mimetic in the active site pocket. The initial structure used in simulations, obtained from Suárez and Díaz.,^23^ is shown. The substrate is coloured in orange, with nitrogen atoms and oxygen atoms coloured blue and red, respectively. M^pro^ side chains coloured in white are those that interact directly with the substrate. Acetyl and N-methyl groups capping the N- and C-termini are omitted for clarity. (D) Metrics used to determine equilibration in the apo dimer simulations. Based on root mean square deviation (RMSD), radius of gyration (R_gyr_), number of hydrogen bonds, and solvent accessible surface area (SASA), the first 0.5 µs was excluded as equilibration. The dashed line delineates the equilibration from the part of the trajectories used for analysis. Shaded regions denote the standard error of the mean from five independent trajectories. (E), (F) Comparison of mean RDCs computed from apo dimer simulations and experimental values.^34^ Points are coloured by the domain to which each residue belongs. The root-mean-square deviation (RMSD) between experimental values and RDCs computed from simulation data are shown on each plot. The Pearson correlation, r, for ^1^D_NH_ and ^2^D_C’H_ RDC values is r = 0.964 ± 0.006 and 0.941 ± 0.009, respectively.

To evaluate the accuracy of our simulations, we computed residual dipolar couplings (RDCs) using the M^pro^ apo dimer simulations. We then compared the computed RDCs to those obtained using NMR spectroscopy by Robertson et al.^34^ (Figure 1E and F). We observe relative agreement between the two, as shown through a mean Q-factor of 0.398 ± 0.007 comparing ^1^D_NH_ and ^2^D_C’H_ RDC values. To put this comparison into context, we note that Robertson et al. report Q-factors of 0.2-0.53 between their measured RDCs and RDCs computed using X-ray crystal structures of M^pro^.^34^ Q-factors of 0.2 to 0.7 have been reported when comparing simulated ensembles using multiple force fields and experimental RDCs in a study evaluating the accuracy of MD simulations.^35^ Differences between experimental RDCs and RDCs predicted from the simulation ensembles may stem from using a C145A active site mutant and an alignment medium in the NMR experiments,^34^ as well as limitations in simulation accuracy, the use of virtual sites, and conformational sampling. Based on this comparison to the available NMR data for M^pro^, as well as additional qualitative comparisons to other structural and biophysical studies described below, we proceed with the analysis of the conformational ensembles of M^pro^ in different conditions.

### 2.2 The Flexibility and States of the Two Active Site Loops

Surrounding the active site of each protomer are two loops, the upper loop (residues 44-54) and the lower loop (residues 184-194), with loop boundaries defined consistently with previous studies.^23,25^ Mutations in both loops occur in SARS-CoV-2 variants of concern.^12^ These mutations increase (L50F and E47K) or decrease (E47N) the enzyme’s activity due to changes in charge distribution and the orientations of side chains in the substrate binding site.^12^ Mutations in both the upper (L50F) and lower loops (V186V, A193P) confer resistance to nirmatrelvir,^13^ suggesting the importance of these loops to the function of M^pro^ and the utility of studying their structure and dynamics in detail. An additional loop of interest due to its interactions with bound ligands comprises residues 162-174; we refer to this loop as the catalytic loop (Figure 1A).

Both the upper and lower loops exhibit high flexibility, as evidenced by their higher root-mean-square fluctuation (RMSF) compared to the rest of the protein (Figure 2C and 3B). This observed flexibility is consistent with other MD simulation studies of M^pro^ ^15,25,26^ and is likely crucial in accommodating different types of residues in the active site. ^4^ Increased flexibility in the upper and lower loops is not evident in crystallographic B-factors (Figure S1). However, both loops are significantly more flexible in solution simulations compared to what would be expected based on the B-factors. An analysis of over 200 crystal structures of M^pro^ reveals that each loop adopts a distinct orientation corresponding to a partially closed conformation in all of these structures (with the upper loop in a closed state and the lower loop in an intermediate state, see section “Substrate Binding Favours Intermediate and Closed Loop States” below), regardless of whether an inhibitor is bound (Figure S2). Furthermore, an analysis of crystal contacts in crystal structures of apo M^pro^ (Figure S1B) indicates that both the upper and lower loops have many crystal contacts. These contacts are likely stabilizing more compact conformations of both loops. The crystallographic structures may, therefore, provide a limited description of the flexibility of the upper and lower loops. These observations are comparable to HIV-1 protease, which similarly exhibits closed or semi-open active site loops in crystal structures and heterogeneous loop conformations in solution.^19^

**Figure 2:**
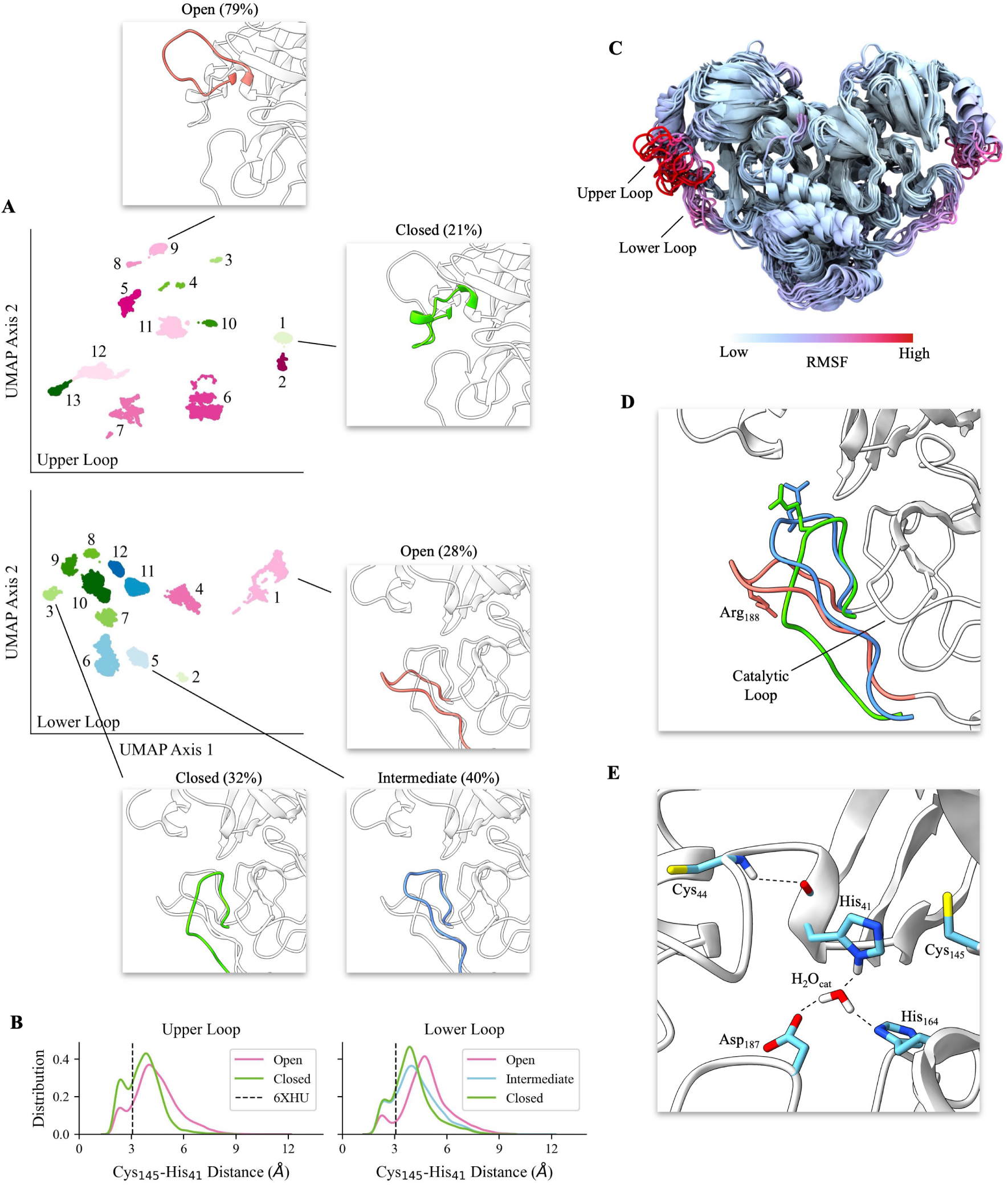
Flexibility and conformational states of the upper and lower loops in the apo dimer. (A) UMAP projections and HDBSCAN clusters of the upper and lower loops, shown in the upper and lower panels, respectively. Numbers within the UMAP plots signify each cluster. Clusters are coloured individually. Pink shades denote open states (upper loop: clusters 2, 5, 6, 7, 8, 9, 11, and 12 and lower loop: clusters 1 and 4). Green shades denote closed states (upper loop: clusters 1, 3, 4, 10, and 13 and lower loop: clusters 2, 3, 7, 8, 9, and 10). Blue indicates an intermediate state (lower loop: clusters 5, 6, 11, and 12). Representative structures and the population of each state are provided in insets. (B) Histograms of the catalytic dyad distance (distance between the Cys_145_ thiol hydrogen and the His_41_ imidazole nitrogen) in the different conformational states of the upper (left panel) and lower (right panel) loops. This distance in the crystal structure (PDB 6XHU) is indicated as a dashed line in both panels. (C) Apo dimer coloured by RMSF with a range of 0 (low) to 6 Å (high). Fifteen structures from the simulation are shown (selected from the trajectory at evenly spaced intervals). The data is shown as a line plot in Figure 3B. (D) Representative structures of open, intermediate, and closed lower loop states are coloured in pink, blue, and green, respectively, with the side chain of Arg_188_ shown. (E) View of the active site taken from the apo dimer simulation with both loops in a closed state. Key hydrogen bonds stabilizing the closed state of both loops are shown as dashed lines. Cys_145_ and His_41_ are the catalytic dyad residues, Asp_187_ and His_41_ are catalytic helper residues, and Cys_44_ and His_41_ form a hydrogen bond that orients His_41_ towards Cys_145_.

**Figure 3:**
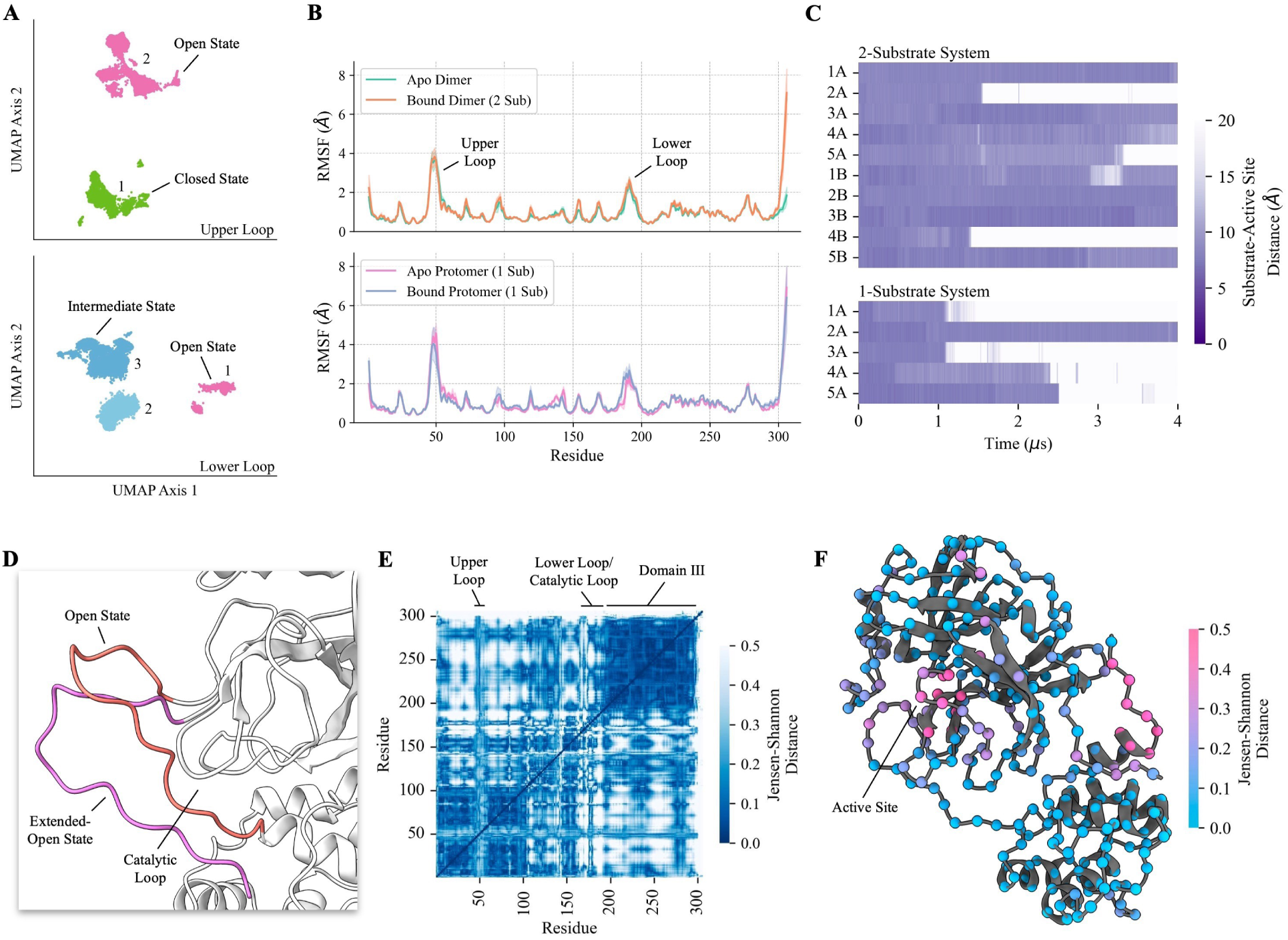
Effects of substrate binding on M^pro^. (A) UMAP projections and HDBSCAN clusters of M^pro^ upper and lower loops in the two-substrate system. Numbers within the UMAP plots label each cluster. Pink, blue, and green shades denote open, intermediate, and closed loop states, respectively. Shades of each colour are assigned arbitrarily. Populations of each state are provided in Table 2. (B) Mean RMSF per residue for the apo dimer and two-substrate system (top) and the bound and apo protomers of the one-substrate system (bottom). Shaded regions denote the standard error of the mean. (C) Distance between the substrate and active site (defined as the distance between Cys_145_ and the closest substrate residue) over the length of the simulations for the two- and one-substrate systems, numbered by trajectory and protomer (e.g. 1A is protomer A of the 1st trajectory). Purple regions indicate that the substrate is in the active site, while white regions show when it has dissociated. (D) Representative structures illustrating the open and extended-open states of the lower loop. (E) Jensen-Shannon distance (JSD) between distributions of C*α* distances, comparing the apo dimer and two-substrate ensembles. Pairs of residues with low JSD (blue) sample similar states between systems, whereas high JSD (white) describes significant ensemble differences. (F) The M^pro^ monomer coloured by the JSD between the apo dimer and two-substrate systems using backbone dihedral angles as features. The spheres represent the C*α* of each residue and are coloured by JSD.

To learn more about the structure and dynamics of the active site loops, we carried out dimensionality reduction using uniform manifold approximation and projection (UMAP),^36^ a general nonlinear reduction technique previously used to study protein folding intermediates and loop motions.^37,38^ We used UMAP because of its scalability with large datasets, increased computational efficiency, better preservation of global structure, and ability to process non-linear data compared to other dimensionality reduction methods.^36^ UMAP constructs a high-dimensional graph representation of a dataset before optimizing a lower-dimensional graph while conserving as much topological similarity as possible. After dimensionality reduction, we used hierarchical density-based spatial clustering of applications with noise (HDBSCAN), whose main advantage over other algorithms is its ability to detect clusters of varying densities.^39^

Because both the upper and lower loops exhibit high flexibility (Figure 2C), we wanted to obtain a quantitative description of the structurally heterogeneous ensembles of each loop. For this purpose, we used UMAP and HBDSCAN to cluster the structural ensemble of the region surrounding the active site. The UMAP projection of the apo dimer’s upper loop yields 13 clusters (Figure 2A), each of which corresponds to a closed or open state of this loop. The clusters are categorized as closed or open based on the distance between the loop tip and residue Pro_9_ in the N-terminus of the protein, which is used as a reference point due to its stability in the structure (see methods, section “Dimensionality Reduction and Clustering”). The loop behaves similarly in all open state clusters — it is free to move around in space with high flexibility. A plot of free energy as a function of the distance between the loop tip and Pro_9_ reveals that conformations near the boundary between the open and closed states are the most energetically favorable, with a moderate preference towards open states (Figure S3A). This preference is further supported by the high population of the open state (79 ± 8%) compared to the closed state (21 ± 8%) (Table 2).

**Table 2:** Summary of the populations of the upper and lower loop states based on UMAP and clustering with HDBSCAN. The number in brackets indicates the standard error of the mean for each population (based on n = 5 independent simulations). Sub refers to substrate in the description of each system.

| System | Loop | Open % | Intermediate % | Closed % |
| --- | --- | --- | --- | --- |
| Apo Dimer | Upper | 79 (8) | - | 21 (8) |
| Apo Protomer from Dimer (1 Sub) |  | 90 (10) | - | 10 (10) |
| Apo Monomer |  | 100 (0) | - | 0 (0) |
| Bound Dimer (2 Sub) |  | 56 (14) | - | 44 (14) |
| Bound Protomer from Dimer (1 Sub) |  | 62 (15) | - | 38 (15) |
| Apo Dimer | Lower | 28 (13) | 40 (11) | 32 (11) |
| Apo Protomer from Dimer (1 Sub) |  | 6 (6) | 44 (14) | 50 (16) |
| Apo Monomer |  | 75 (11) | 25 (11) | 0 (0) |
| Bound Dimer (2 Sub) |  | 13 (8) | 87 (8) | 0 (0) |
| Bound Protomer from Dimer (1 Sub) |  | 30 (10) | 70 (10) | 0 (0) |

Applying UMAP and HDBSCAN on the apo dimer’s lower loop reveals 12 clusters corresponding to open, closed, and intermediate states (Figure 2A, lower panel). The free energy profile reveals that the intermediate and closed states are more energetically favorable than the open state, with the lowest energy conformations occurring near the boundary between the intermediate and closed states (Figure S3B). In the closed states, the catalytic loop shifts downwards, enabling the lower loop to curl inward into the active site. This allows for a more compact conformation in solution compared to what is observed in crystal structures. The lower loop shifts further away from the catalytic loop as it moves from closed to intermediate to open (Figure 2D), suggesting that the state of the lower loop may play a role in substrate binding and catalysis.

The flexibility of the two loops observed in our simulations is consistent with the work of Gossen et al., who described the flexibility and conformational changes of M^pro^’s active site through Markov state model analysis.^14^ While they did not identify distinct loop states as we do in the present study, they noted that the volume of the binding site is affected by the flexibility of the loops, suggesting that the movement of loops, and possibly different loop states, may contribute to creating more or less favourable binding conformations. Lee et al. have obtained several crystallographic structures of M^pro^ bound to different substrates, which reveal slight structural differences in the upper and lower loops depending on the bound substrate. These findings provide strong evidence for loop flexibility, which is likely functionally important in accommodating various substrates.^3^

### 2.3 Catalytic Residues are Stabilized in Closed Loop States

Next, we investigated if the conformational states of the upper and lower loops correlate with the active site’s conformation. The distance between the catalytic dyad residues (Cys_145_ and His_41_) in the apo dimer is ≤ 4.0 Å on average when either loop is in a closed state (Table 3). The Cys_145_–His_41_ distance is also within error of the closed state distance when the lower loop is in an intermediate state, suggesting some similarity between these two states. In contrast, the Cys_145_–His_41_ distance rises to an average range of 4.4-4.8 Å when either loop is in an open state, with wider distributions of this distance compared to closed and intermediate states (Figure 2B). This variation in the distance between the catalytic dyad residues is likely to affect the catalytic efficiency. Additionally, when the upper loop is closed, the side chain of His_41_ and the Cys_145_ thiol become more rigid compared to when the loop is open, based on an analysis of the RMSF of these residues (Figure S4). This increased rigidity may contribute to the reduced variance and shorter distance between the two catalytic dyad residues. The cause of the increased rigidity may be the closer proximity of the backbone nitrogen of Cys_44_ and the carbonyl oxygen of His_41_ in the closed state (Table 3), where a hydrogen bond is typically present (Figure 2E). The shorter distance may strengthen the hydrogen bond, stabilizing the His_41_ backbone, which in turn stabilizes both its side chain and the interacting Cys_145_ residue. Taken together, these results indicate that Cys_145_ and His_41_ are more likely to be close enough for proton transfer to occur when at least one of the loops is in a closed or intermediate state, suggesting that closed and intermediate states are more favourable for catalysis.

**Table 3:** Catalytic site distances in different states of the upper and lower loops. Values in brackets denote the standard error of the mean for each distance. Residue Cys_44_ is in the upper loop and residue Asp_187_ is in the lower loop. For clarity, only distances that are relevant for comparison between different loop states are shown, and distances that are not relevant for comparison are indicated by dashes.

| Loop | Loop State | Cys<sub>145</sub>-His<sub>41</sub> | Asp<sub>187</sub>-His<sub>41</sub> | Cys<sub>44</sub>-His<sub>41</sub> |
| --- | --- | --- | --- | --- |
| **Mean Distance (Å) in the Apo Dimer** |  |  |  |  |
| Upper | Open | 4.4 (0.1) | - | 3.1 (0.3) |
|  | Closed | 3.7 (0.2) | - | 2.6 (0.2) |
| Lower | Open | 4.8 (0.1) | 7.5 (0.8) | - |
|  | Intermediate | 4.2 (0.3) | 5.1 (0.1) | - |
|  | Closed | 4.0 (0.1) | 4.90 (0.09) | - |
| **Mean Distance (Å) in the Bound Dimer** |  |  |  |  |
| Upper | Open | 4.3 (0.2) | - | 3.4 (0.3) |
|  | Closed | 3.7 (0.2) | - | 2.35 (0.01) |
| Lower | Open | 5.6 (0.3) | 8.8 (0.4) | - |
|  | Intermediate | 3.8 (0.1) | 6.9 (0.1) | - |
|  | Closed | - | - | - |

We observe additional effects on the conformation of the active site in different states of the upper and lower loops. Similarly to the shorter Cys_145_–His_41_ distance in closed loop states, a shorter distance is also observed between Asp_187_ and His_41_ when the lower loop is in an intermediate or closed state (Table 3). The interaction between these two residues is thought to be mediated by a water molecule (H_2_O_cat_, Figure 2E) that plays an essential role in catalysis.^4,40,41^ It stabilizes the positive charge of protonated His_41_ through electrostatic interactions with the negatively charged Asp_187_ during cleavage.^3,33^ In crystal structures,^4,40,41^ as well as in our simulations, this water molecule forms hydrogen bonds with His_41_, His_164_, and Asp_187_. When the lower loop is in an intermediate or closed state, the mean Asp_187_-His_41_ distance is 5.1 ± 0.1 or 4.90 ± 0.09 Å, respectively, and water-bridged interactions between His_41_, His_164_, and Asp_187_ are formed 64 ± 16 and 85 ± 9% of the time, respectively. The open state has a water molecule bridging these three residues only 7 ± 3% of the time, as the mean Asp_187_-His_41_ distance increases to 7.5 ± 0.8 when open. These results provide additional evidence that the closed and intermediate lower loop states may be more conducive to catalysis than the open state.

Next, we wanted to determine the extent to which the state of one loop influences the state of the opposite loop. We observe that when the upper loop is closed, the lower loop is in an intermediate or closed state 17.5% more often than average. When the upper loop is open, the lower loop is open 4.5% more often than average. Additionally, when the lower loop is open, the upper loop is found to be open 14.5% more than average (Table S1). These results may indicate a cooperative anchoring of closed states. This cooperativity may arise from the orientation of the Arg_188_ side chain on the lower loop in different loop states. The positively-charged Arg side chain is oriented upwards toward the upper loop in closed and intermediate states, allowing it to form inter-loop interactions that bridge the two loops. In contrast, the Arg_188_ side chain often faces downwards in an open state (Figure 2D), resulting in fewer interactions with the upper loop. The reorientation of this side chain is reflected in time series plots of the Arg_188_-Pro_9_ distance and *ψ* angle (Figure S5). Closer Arg_188_-Pro_9_ distances generally exhibit *ψ* angles near 180*^◦^*, while larger distances show a shift of this angle toward 0*^◦^*. While Arg_188_ is moderately conserved in coronaviruses (Figure S6), other residues in the lower loop may play similar roles in stabilizing closed loop states when the Arg is not present, such as the lysine residue in the same position in M^pro^ of MERS-CoV. As the closure of either loop stabilizes catalytic residues (Table 3), cooperativity in the closure of loops is likely key to efficient cleavage of the substrate. Inhibitors designed to restrict the closure of one loop may indirectly limit the closure of the other, providing a unique mechanism for inhibiting the protease.

### 2.4 Substrate Binding Favours Intermediate and Closed Loop States

To determine the effects of substrate binding on the loops, we carried out the same dimensionality reduction and clustering analysis on the two-substrate system (Figure 3A) and the one-substrate system (Figure S7). When comparing the two-substrate system to the apo dimer system, we find that substrate binding increases the probability of the intermediate state in the lower loop while decreasing the probability of the open state (Table 2). Due to steric hindrance, the lower loop cannot sample a maximally closed state when a substrate is present in the active site. This is reflected in the free energy profile of the lower loop upon binding, where the closed state becomes highly unfavorable, and the free energy minimum shifts to the intermediate state (Figure S3D). However, the mean distance between His_41_ and Cys_145_ in intermediate and closed states are the same within error, suggesting that either state may be suitable for catalysis (Table 3). Upon substrate binding, the closed state in the upper loop becomes more energetically favorable than in the apo dimer (Figure S3C). This leads to an increased frequency of the closed state, further reducing the His_41_-Cys_145_ distance and rigidifying the catalytic dyad through stronger Cys_44_-His_41_ interactions, promoting configurations suitable for catalysis.

Upon substrate binding, several regions in the dimer exhibit differences in structure. We assessed these changes in the structural ensemble quantitatively using the Jensen-Shannon distance (JSD) to compare the apo and bound ensembles in terms of inter-residue distances (Figure 3E) and backbone dihedral angles (Figure 3F). The JSD is a symmetrized measure of the similarity of probability distributions, with values closer to zero indicating similarity and values closer to one indicating dissimilarity. ^42,43^ Computing the JSD between distributions of C*α* distances and distributions of dihedral angles reveals that the most significant differences upon binding occur in the active site loops and the catalytic loop, with moderate changes occurring at the C-terminus and the dimerization interface (Figure 3E-F). In contrast, Domain III exhibits only minor structural differences between the apo and bound ensembles. The fact that the most prominent differences occur in the vicinity of the active site is expected because of the increased frequency of closed/intermediate loop states in the bound system and the interactions between the substrate and active site.

The observed states of M^pro^’s active site loops are analogous to other enzymes with flexible active site loops that have distinct apo and bound conformations. For example, Du et al. found that fructose-1,6-bisphosphate aldolase/phosphatase has three active site loops that changed conformation when either a substrate, reaction intermediate, or product is bound.^44^ They observed that the loops in the aldolase/phosphatase reorient to bring key side chains and Mg^2+^ cofactors into place. This functional role is comparable to the lower loop of M^pro^, which brings Asp_187_ closer to the catalytic dyad in a water-bridged interaction. The behaviour of the active site loops of M^pro^ also resembles HIV-1 protease, which has different frequencies of loop conformational states based on the presence or absence of a substrate.^18,19^ The loop flexibility in HIV-1 protease has been proposed to have a functional role: product release may be facilitated by the higher conformational entropy of the loops surrounding the active site in the apo compared to the bound state.^19^ A similar mechanism may be important in the function of M^pro^.

### 2.5 Substrate Binding in One Protomer Correlates with a Bound-Like Conformation in the Apo Partner

To determine the effect of substrate binding on the rest of the protein, we conducted a detailed analysis of the one-substrate system. We used UMAP and HDBSCAN to analyze both protomers in the one-substrate system (Figure S7), following the same approach we used to analyze the loop states in the apo dimer (Figure 2). We find that the apo protomer’s upper loop samples open and closed states at frequencies comparable to the apo dimer (Table 2). The lower loop, however, samples the intermediate and closed states significantly more than the open state, comparable to the two-substrate bound dimer despite the absence of substrate (Table 2). The similarity in loop state frequencies between the two-substrate system and the apo protomer from the one-substrate system suggests that binding in one protomer induces a bound-like ensemble in its apo partner.

In addition to the differences in loop states, the one- and two-substrate systems also differ in their retention of the substrate. The substrate dissociates from the active site within 4 µs in 80% of the trajectories of the one-substrate system, compared to only 30% of the two-substrate system (Figure 3C). As we started both simulation systems from the same initial structure and under the same starting conditions (except for the presence of the substrate in one of the protomers), we do not expect the observed difference in dissociation to be a simulation artifact. As shown above, binding in one protomer in the one-substrate system contributes to the increased sampling of intermediate/closed states of its partner’s active site. However, while the bound protomer contributes to the closing of the apo protomer’s active site, the apo protomer does not reciprocate this effect in the bound protomer. This can be seen in the increased frequency of the open lower loop state in the bound protomer (30 ± 10%) compared to the apo protomer (6 ± 6%), which is likely what causes the quicker substrate dissociation in the one-substrate system (Figure 3C). Overall, the observed dependence of the conformational ensemble on the presence or absence of substrate in the opposite protomer suggests that there is allosteric communication between protomers.

### 2.6 Dynamic Network Analysis

The similarity of the behaviour of the lower loop in the apo protomer and its bound partner in the one-substrate system (Table 2) is suggestive of allosteric communication between protomers. To describe potential allosteric pathways connecting the two protomers and associated correlated motions, we used dynamic network analysis (DNA). DNA uses cross-correlation to reduce trajectories into a “nodes and edges” representation, which can then be analyzed as a network.^45,46^ In this analysis, each residue is classified as a node, and nodes are linked to each other through edges, each with an associated correlation coefficient. The network can then be partitioned into “communities” of dense connections, and optimal paths can be determined by computing the shortest pathways between all nodes.^46,47^

The cross-correlation coefficients computed from the simulations of the one-substrate dimer (Figure 4A) reveal the correlated motion of residues within each domain, consistent with other studies using cross-correlation^30,32^ or linear mutual information.^27^ The fact that there is moderate correlation between the two active sites (box 1 in Figure 4A) provides further support for allosteric communication occurring between the two protomers. The motion of the N-terminus of one protomer is correlated with the opposite protomer’s active site (box 2 in Figure 4A), consistent with the fact that the N-terminus contributes to shaping the active site of the opposite protomer. ^7^ Using these correlation coefficients, the system can be decomposed into nine communities of high information transfer (Figure 4B), which match well with the defined boundaries of the three domains of M^pro^ (Figure 1A). The system can also be represented as a network with edges weighted by their “betweenness” (Figure 4E), which is defined as the number of shortest paths from all nodes to all others that pass through that edge. In other words, a high betweenness (represented as a thicker red line) corresponds to an avenue of high information transfer.^45^ From this network view (Figure 4E), it can be seen that much of the protein’s information transfer travels directly through the middle of the protein, strongly suggesting that the dimer interface is important in allosteric communication.

**Figure 4:**
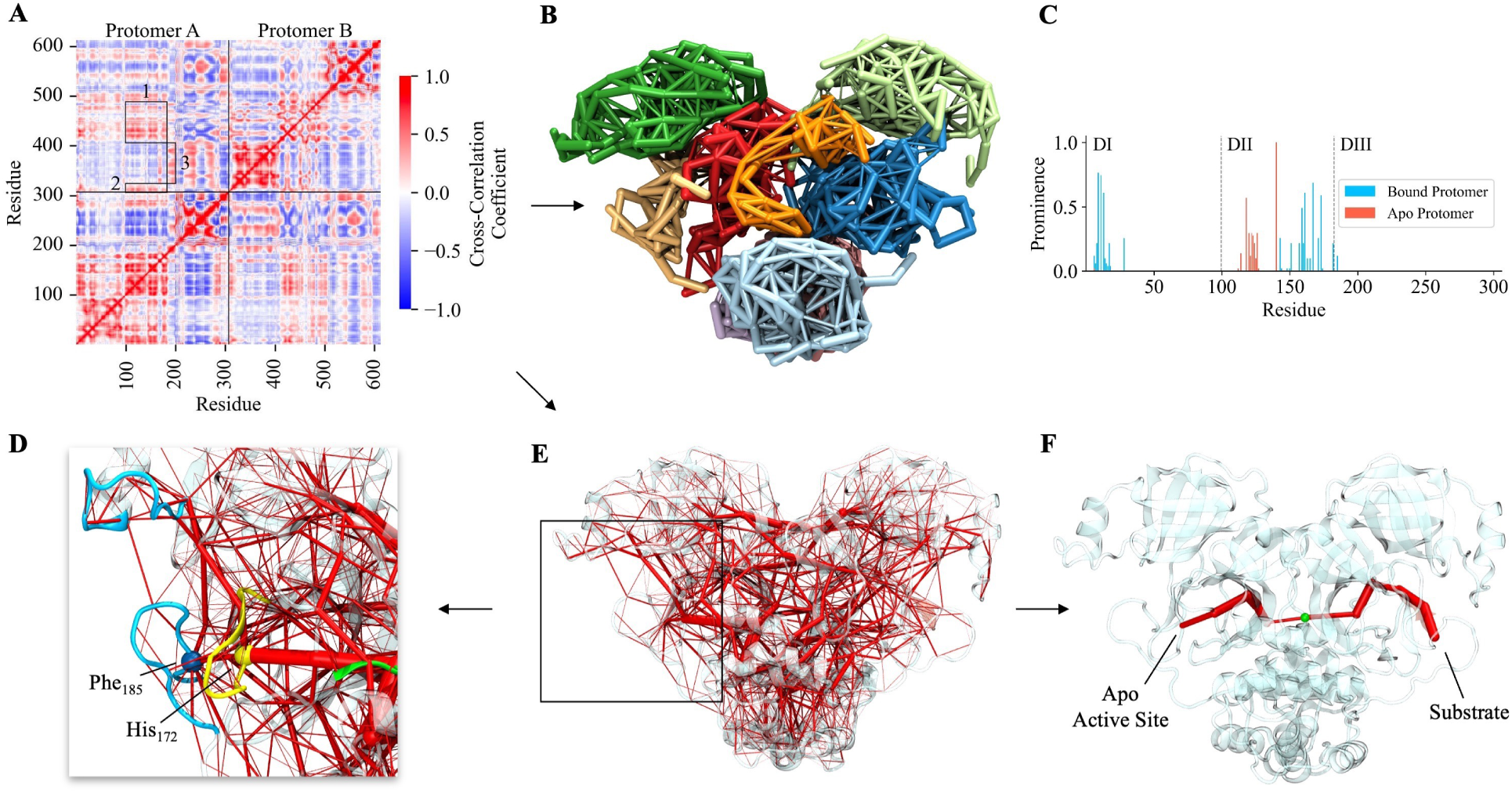
Dynamic network analysis of the one-substrate system. (A) Cross-correlation coefficient matrix. Values close to 1, −1, and 0 show correlation, anti-correlation, and no correlation, respectively. Boxed areas show a notable correlation between protomers, which are split by horizontal and vertical black lines. Box 1 shows correlation between Domain IIs, box 2 between the N-terminus and the opposing active site, and box 3 the lower loop and Domain I of the opposite protomer. (B) M^pro^ clustered into nine communities (note that some of the communities are on the opposite side and not visible in this orientation of the dimer). (C) Prominence of each residue, i.e. how often each residue is involved in the 50 shortest paths from one active site to the other. Boundaries between the three domains are indicated by dashed lines. (D) Enlarged view of the apo protomer’s active site. The loops are coloured blue, the N-terminus green, and the catalytic loop yellow. Phe_185_ and His_172_ serve as hub nodes (spheres) with several connections to other parts of the active site. (E) Network representation of M^pro^ with edges weighted by betweenness, defined as the number of shortest paths from all nodes to all others that pass through that edge; the thicker the edge, the higher the betweenness. (F) Shortest path from the substrate to the apo protomer’s active site. The optimal (shortest) path (red) travels through Met_6_ in the N-terminus (green sphere).

In dynamic network analysis, the shortest distance between two sites corresponds to the optimal path of communication, as shorter distances result in enhanced allostery compared to suboptimal paths.^46^ Analysis of the M^pro^ dimer simulations reveals that the shortest pathway between the active sites travels through Domain II of both protomers, which are bridged by one protomer’s N-terminal region (specifically, through residue Met_6_) (Figure 4F). Computing the 50 shortest paths between the active sites shows that pathways involving several residues in the N-terminal region and Domain II are involved in communication (Figure 4C). The shortest paths consistently pass directly through the center of the protein, including the dimer interface, and through the N-terminus of either protomer (Figure S8), suggesting that any one of these regions may be a target for obstructing allosteric communication. If this communication between protomers is essential to the function of M^pro^, allosteric inhibitors that bind at the dimer interface and disrupt this communication may disrupt activity and provide an alternative to competitive inhibition. Such an approach to allosteric modulation of M^pro^ by targeting its dimer interface has been suggested in prior studies.^6,31,48,49^ For instance, Douangamath et al. identified three fragments that bind M^pro^ at the interface near the N-terminus,^48^ while Cantrelle et al. identified an allosteric hot spot at the interface using NMR spectroscopy.^49^ Douangamath et al. identified multiple non-covalent binders that bind at the dimer interface,^48^ one of which has been shown to promote dissociation of the dimer. ^6^ In our analysis, we find that the N-terminus has a path of high betweenness to His_172_, which acts as a hub node (Figure 4D-F). His_172_ is closely connected to both loops, as well as the active site (Figure 4D). His_172_’s high betweenness with Phe_185_, a hub node on the lower loop, suggests that it may play a role in the conformational change of the loops upon binding. The upper loop is also closely connected to the lower loop through short pathways from either His_172_ or Phe_185_. The results from dynamic network analysis are supported by prior studies demonstrating the functional importance of His_172_ and Phe_185_. In particular, protonation of His_172_ has been shown to result in a collapse of the dimer interface^29,30^ and a partial collapse in the catalytic site, ^20,29^ suggesting its importance in the structure and allostery of M^pro^. Phe_185_ was identified as a mutational coldspot through an analysis of missense mutations reported in the Global Initiative on Sharing All Influenza Data (GISAID).^50^ The lack of mutations in key catalytic residues suggests that coldspots are likely to be functionally important, which agrees with our identification of Phe_185_ as a functionally important residue due to its role as a communication hub. Additionally, Chen et al. identified a mutation at this site, F185S, that exhibited a 30-fold decrease in activity, providing further evidence of the importance of Phe_185_.^12^ Furthermore, both His_172_ and Phe_185_ are highly conserved among coronaviruses (Figure S6).

N-terminal residues are involved in six of the top ten edges of highest betweenness between protomers (Table S2), signifying the importance of the N-terminus in interprotomer communication. N-terminal residues also populate the list of residues with the most connections between protomers (Table S3). Specifically, Arg_4_ acts as a communication hub, with several total edges and edges of high betweenness connecting the two active sites (Table S2 and Table S3). Arg_4_ has also been reported to play a role in dimerization, forming a salt bridge with Glu_290_ on Domain III of the opposite protomer^7^ and providing an allosteric path between Domain II, the opposite N-terminus, and Domain III. El Ahdab et al. also showed that the distance between the catalytic dyad may be allosterically linked to the distance between Arg_4_ and Glu_290_.^30^

Previous computational studies have reported allostery in M^pro^.^32,48,51–53^ Elastic network modelling by Dubanevics et al. revealed allostery between active sites, which is consistent with the results of our network analysis.^52^ Sencanski et al. and Verma et al. discovered allosteric inhibition pockets and noted that inhibitory constants at these sites were significantly stronger than competitive inhibition pockets, with the added benefit that inhibitors need not compete for the active site.^54,55^ Campitelli et al. used a dynamic coupling index to quantify allostery between the catalytic Cys_145_-His_41_ dyad and Glu_55_, Ile_59_, and Arg_60_ residues on the opposite protomer, all of which are located in the upper loop.^53^ Lastly, Flynn et al. provide evidence of a chain of mutationally-sensitive residues that connect the active site to the dimerization interface, and each of these residues is crucial to protein function.^56^ Using the one-substrate system, we have shown that M^pro^’s active sites are allosterically connected, and using DNA, we find that the optimal allosteric path travels through the middle of Domain II of both protomers in a pathway bridged by the N-terminus. The N-terminus is a key part of the allosteric pathways that facilitate communication with the partner protomer’s active site, dimerization domain, and both the upper and lower loops. Furthermore, we have identified three key hub residues in the allosteric network (His_172_, Phe_185_, and Arg_4_), which play functionally important roles due to their connections to the dimer interface or to one or both active sites. These residues are likely to serve as promising allosteric drug targets.

### 2.7 Stability of the M^pro^ Monomer

To expand on the importance of dimerization in the allosteric communication and the dynamics of M^pro^, we evaluated the differences between monomeric and dimeric M^pro^ assemblies by comparing our monomer and dimer simulations. Based on a comparison of the RMSF profiles, the monomer is significantly more flexible than the dimer throughout the entire protein, but especially in the N-terminus, the dimer interface (including residues 120-170), and the lower loop (Figure 5A). This difference in flexibility is consistent with other studies,^25,26^ and is likely because these are the regions of M^pro^ that have stabilizing interactions in the dimer.

**Figure 5:**
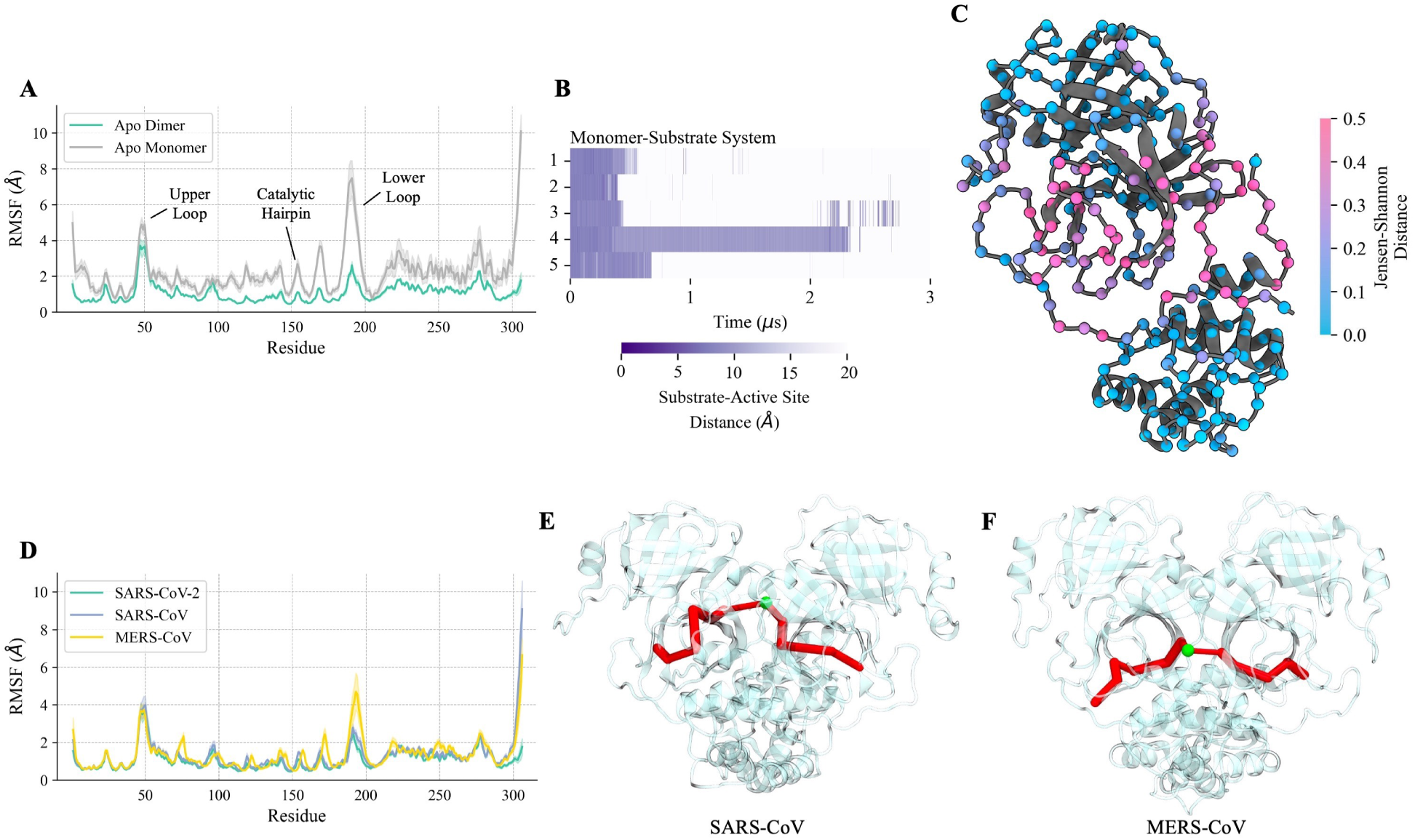
A comparison of the SARS-CoV-2 apo dimer with the apo monomer, as well as with the SARS-CoV and MERS-CoV dimers. (A) Mean RMSF per residue for the SARS-CoV-2 apo dimer and apo monomer. Shaded regions denote standard error of the mean. (B) Distance between the substrate and active site (defined as the distance between Cys_145_ and the closest substrate residue) over the length of the simulations for the SARS-CoV-2 monomer-substrate complex, numbered by trajectory. Purple regions indicate that the substrate is in the active site, while white regions show when it has dissociated. (C) The M^pro^ monomer coloured by the JSD between the SARS-CoV-2 apo dimer and apo monomer using backbone dihedral angles as features. The spheres represent the C*α* of each residue and are coloured by JSD. The largest differences between the apo monomer and apo dimer ensembles occur in Domain II. (D) Mean RMSF per residue for SARS-CoV-2, SARS-CoV, and MERS-CoV apo M^pro^ dimers. Shaded regions denote standard error of the mean. (E), (F) Shortest path between active sites in SARS-CoV and MERS-CoV M^pro^s. The optimal (shortest) path (red) travels through the N-terminus (green sphere).

UMAP projections and HDBSCAN clustering of the apo monomer simulations indicate that both the upper and lower loops of the monomer adopt open states more frequently than in the dimer (Figure S9A, Table 2). In the apo monomer, the lower loop no longer samples the closed state. Analysis of the structural ensembles of the monomer and dimer suggests that the major structural difference preventing the lower loop from adopting the closed state is the absence of a hydrogen bond between Glu_166_ of one protomer and Ser_1_ of the opposite protomer. This interaction, which cannot occur in the monomer, stabilizes the catalytic loop in the dimer, and its absence results in a more flexible catalytic loop and a change in the structure of the active site, as is evident in the JSD analysis (Figure 5C). The increased flexibility of the catalytic loop results in fewer stabilizing hydrophobic interactions with the lower loop, leading to frequent sampling of an “extended-open” conformation in which the lower loop is dislodged from the catalytic loop (Figure 3D). The frequent sampling of the open state by the lower loop causes dissociation of the substrate within 1 µs in all monomer-substrate complex simulations (Figure 5B). In contrast, the dimer-substrate complex exhibits less dissociation (Figure 3C). Taken together, these results suggest that the rigidity of the lower loop upon dimerization is crucial for retaining the substrate, which may contribute to the experimentally observed inactivity of the M^pro^ monomer.^5,6^ Our results are also consistent with a mass spectrometry study by El-Baba et al., which found that the monomer is not only inactive but also does not bind substrate with high affinity.^6^

Dimerization of M^pro^ imparts stability to the active site, as the partner protomer’s N-terminus helps form the catalytic pocket through hydrogen bonding.^5,57^ In our simulations, we also see a positional shift in Domain III upon dimerization (Figure S10A), as shown through high JSDs between Domain III and the rest of the protein when comparing the dimer and monomer ensembles (Figure S10B). In related coronaviruses (HCoV 229E and TGEV), Domain III has been shown to hold the lower loop in a catalytically competent position (in TGEV),^58^ and truncation of this domain in *in vitro* experiments reduced the activity of the protease (in HCoV 229E, TGEV).^58,59^ Truncation of Domain III in SARS-CoV-2 M^pro^ was similarly found to decrease activity due to a shift in the monomer-dimer equilibrium towards the monomer.^60,61^ The shift seen in Domain III upon dimerization likely reorients the domain into a more catalytically favourable position that stabilizes the lower loop. These factors explain the instability of the monomer’s active site and the dissociation of the substrate observed in simulations of the monomer.

### 2.8 SARS-CoV and MERS-CoV M^pro^ dynamics and allostery are similar to SARS-CoV-2

To determine the extent to which our findings can be generalized to M^pro^ of other coronaviruses, we carried out MD simulations of SARS-CoV and MERS-CoV M^pro^ dimers in the apo state. Both coronavirus M^pro^s show strong similarities to SARS-CoV-2 M^pro^ with respect to dynamics and allostery. The RMSF profiles indicate similar flexibility throughout most of the protein (Figure 5D), which is unsurprising considering their high sequence similarity (96.1% and 50.8% for SARS-CoV and MERS-CoV M^pro^ compared to SARS-CoV-2 M^pro^; sequence alignment provided in Figure S6). As we observe for SARS-CoV-2 M^pro^, the upper and lower loops exhibit high flexibility.

Network analyses of SARS-CoV and MERS-CoV M^pro^s (Figure S11) show comparable outcomes to SARS-CoV-2, including similar allosteric pathways between active sites through Domain II of both protomers (Figure 5E and F). Slight differences in optimal paths may be attributed to sampling distinct regions of conformational space among coronavirus M^pro^s, as demonstrated in a Markov-state modelling study of M^pro^ from all three coronaviruses.^15^ In SARS-CoV-2 M^pro^, hydrophobic interactions between Ala_285_ and Leu_286_ from opposite protomers are known to contribute to the stability of the dimer interface. ^57^ We find edges of high betweenness connecting these two residues in the network obtained using DNA (Figure S12A). In M^pro^ from SARS-CoV, the residues at these positions are Thr_285_ and Ile_286_. Instead of hydrophobic interactions, these residues establish only a few contacts, resulting in a weaker bridge between the two protomers. Mutation of these two residues (along with Ser_284_) to Ala in M^pro^ from SARS-CoV resulted in enhanced dimerization due to tighter packing and increased activity by a factor of 3.6.^62^ The tighter packing may be directly related to the increased activity, as a shorter distance between Arg_4_ and Glu_290_ at the dimerization interface tends to correlate with a shorter catalytic distance through allosteric communication. ^30^ The enhanced packing increases the stability of the N-terminus and the active site, as position 286 (along with 277 and 279) is strongly coupled to the active site, and in SARS-CoV, much of this coupling is lost. ^53^ The significance of having different residues at these sites in different coronaviruses is reinforced by the presence of fewer edges between Thr_285_ and Ile_286_ in our DNA model of the SARS-CoV M^pro^ compared to the multiple edges of high betweenness between Ala_285_ and Leu_286_ in the SARS-CoV-2 model (Figure S12).

Experimental studies on SARS-CoV M^pro^ have shown that alanine point mutations at the dimer interface in Domain II disrupt catalytic activity while retaining the global structure of the dimer.^63,64^ This reduction of activity, despite a lack of observed change in structure, suggests that communication between active sites may play a role in catalysis in SARS-CoV M^pro^. Similarly, mutational screens have revealed several residues at the dimer interface that affect the activity of SARS-CoV-2 M^pro^.^56^ As we have shown that allosteric paths between active sites travel through the dimer interface, mutations of interface residues could disrupt communication between protomers. The dimer interface provides a hot spot for several potential inhibitors, as revealed by both experimental (mass spectrometry, X-ray crystallography) and computational studies (docking, algebraic topology). ^6,32,48,49^ We hypothesize that inhibition at this interface may act similarly to the point mutations in reducing catalytic activity by disrupting interprotomer communication.

## 3 Conclusions

Here, we present a combined 92.5 µs of all-atom simulation data on M^pro^s from multiple coronaviruses in different forms. First, we find that M^pro^’s flexible active site loops sample distinct open, intermediate, and closed conformational states and that closed states of the loops stabilize catalytic residues. We also show that substrate binding increases the frequency of the catalytically favourable closed state. Analysis of loop states provides evidence for allosteric communication between the two protomers. To investigate allosteric transmission in M^pro^, we use dynamic network analysis, finding potential allosteric paths between the active sites in opposite protomers. We observe the instability of the monomer compared to the dimer due to an increased flexibility, which results in its inability to retain the substrate. Finally, we find similarities between M^pro^s from different coronaviruses in terms of loop states, flexibility and allosteric paths, which makes it possible to generalize our findings to M^pro^ from related viruses. The flexibility of the upper and lower loop likely plays a functional role in allowing M^pro^ to bind diverse substrates — not only the 11 cleavage sites in the polyproteins of SARS-CoV-2, but also proteins within the human proteome.^65^ As substrate binding in one protomer is found to be correlated with the behaviour of the other, inhibitors that target allosteric pathways may provide an alternate mode of inhibition.

## 4 Methods

### 4.1 Experimental Design

We carried out extensive all-atom simulations of M^pro^ in several conditions (see Table 1 for a list of simulations). To investigate the effects of dimer against monomer, apo against substrate bound, and between different coronaviruses, we simulated several systems as follows. The crystal structure PDB 6XHU^33^ was used for the apo SARS-CoV-2 M^pro^ monomer and dimer simulations, while a modified version of a crystal structure containing a covalent inhibitor (PDB 6LU7^8^) obtained from Suárez and Diaz^23^ was used for substrate-bound simulations. In all SARS-CoV-2 M^pro^ simulations, lysines, arginines, aspartates, and glutamates were all charged. His_64_, His_163_, His_164_, His_172_, and His_246_ were all protonated at N*ɛ*, while His_41_ and His_80_ were protonated at N*δ*, consistent with a pH of 7.0.^23^ The bound systems contained a pentapeptide substrate mimic in the active site (details of the substrate can be found in SI Section 1.1). SARS-CoV and MERS-CoV M^pro^ simulations used crystal structures PDB 1UJ1^66^ and PDB 5C3N,^67^ respectively, as the starting structure for the simulations. The structure 1UJ1 is among the highest resolution apo structures of SARS-CoV M^pro^, while 5C3N was the only published crystal structure of MERS-CoV M^pro^ at the start of the study. The protonation states of all residues in the SARS-CoV simulations match those in the apo dimer simulations. In the MERS-CoV simulations, His_8_, His_64_, His_71_, His_175_, and His_194_ are all protonated at N*ɛ* while His_41_, His_83_ and His_166_ are protonated at N*δ*, consistent with a pH of 7.0.^23^ All systems studied, as well as the number of independent trajectories and the length of each simulation, are provided in Table 1. Simulations on a µs timescale were chosen based on the simulation timescale of other M^pro^ simulation studies.^22–25,27,28^

### 4.2 MD Simulations

GROMACS 2019.1 was used for all MD simulations.^68^ The system was composed of the M^pro^ monomer or dimer with charged termini in a rhombic dodecahedral box filled with CHARMM-modified TIP3P water molecules^69^ at a concentration of 0.15 M NaCl for a total of about 87000 atoms for the dimer systems and about 63000 atoms for the monomer systems. The protein had a minimum distance of 10 Å to all box edges in the initial structure, and periodic boundary conditions were applied. The CHARMM36m force field^70^ was used, and virtual sites for all hydrogens were employed to allow for an integration timestep of 4 fs. MkVsites^71^ was used to generate bond constraints for termini capping groups. Energy minimization was done prior to production runs using the steepest descent algorithm. Hydrogen bond lengths were constrained using the LINCS algorithm.^72^ Prior to the production run, a restrained simulation was performed for 5 ns with position restraints on all heavy atoms. Short-range electrostatic interactions and van der Waals interactions were calculated with a 0.95 nm cut-off. Long-range electrostatic interactions were calculated using fast smooth particle-mesh Ewald summation with a fourth-order interpolation and a grid spacing of 0.12 nm.^73^ Neighbour searching was done using the Verlet cutoff scheme. Temperature coupling was done using velocity rescaling to maintain a temperature of 298 K for all simulations.^74^ Equilibration simulations were run in the NPT ensemble for 10 ns using Berendsen pressure coupling,^75^ followed by 10 ns using the Parrinello-Rahman barostat.^76^ Production runs were performed in the NPT ensemble at 298 K and 1 bar using the Parrinello-Rahman barostat and a velocity rescaling thermostat.^74,76^

### 4.3 Calculation of Predicted Residual Dipolar Couplings from Simulations

RDCs were computed from the M^pro^ apo dimer simulation ensembles. A total of 17000 structures from the equilibrated portions of the trajectories were used. Alignment tensors were fit to the experimental data from Robertson et al.^34^ using the DC Order Matrix Fitting function (singular value decomposition) from NMRPipe.^77^ We followed the same procedure as Robertson et al. in their comparison between RDCs computed from crystal structures and experimental RDCs. They optimized the experimental alignment tensor accuracy by iteratively restricting the fit to residues 5-115 and 199-272.^34^ We likewise restricted the fit to these two contiguous regions in M^pro^.

Q-factors were computed via NMRPipe^77^ using the following equation:

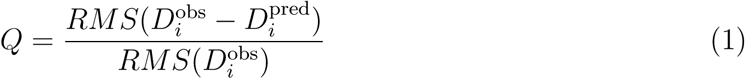

where 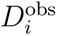 and 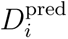 are the observed (from NMR) and predicted (from simulations) RDCs for interaction *i*.^78^

### 4.4 Dimensionality Reduction and Clustering

UMAP^36^ was performed on the upper and lower loops for each system using heavy atom positions as features after alignment of the protein by backbone atoms, excluding the loops and termini. The number of neighbours parameter was chosen to maximize global topological structure in high dimensions over local structure (higher values), while the minimum distance parameter was optimized for clear clusters (lower values). The specific values used were number of neighbours = 400 and minimum distance = 0.1. Clustering was then carried out with HDBSCAN,^39^ where the minimum cluster size and the minimum samples parameters were chosen to maximize relative validity indices (specific values used varied between minimum samples = 20-80 and minimum cluster size = 700-1700). Other choices of features (dihedral angles and backbone interatomic distances) were also investigated, but heavy atom positions provided the clearest categorization of conformational states. Loop states were categorized based on the distance between the loop tip (the C*α* atoms of Leu_50_ on the upper loop and Arg_188_ on the lower loop) and Pro_9_ in the N-terminus, which is used as a reference point due to its stability in the structure (Tables S4-S9). In the upper loop, clusters were determined to be closed if this distance was < 38.5 Å and open if ≥ 38.5 Å. In the lower loop, clusters were determined to be closed if the distance was ≤ 32.5 Å, intermediate if between 32.5 and 34.5 Å, and open if ≥ 34.5 Å. These bins were determined using the same distances in six apo crystal structures (6YB7, 6Y2E, 6WTM, 6M03, 6WQF, and 6XHU) as a guideline, where the upper and lower loops are in closed and intermediate states, respectively. In the crystal structures, these distances are: Leu_50_-Pro_9_ distance = 37.14 ± 0.03 Å and Arg_188_-Pro_9_ distance = 32.73 ± 0.02 Å. Additionally, structures belonging to each cluster were visually inspected to confirm that the assigned loop state was consistent with the position of the loop in simulation frames.

### 4.5 Cross-Correlation and Dynamic Network Analysis

Cross-correlation coefficients were calculated using CARMA.^79^ Pairwise correlations were calculated as follows:

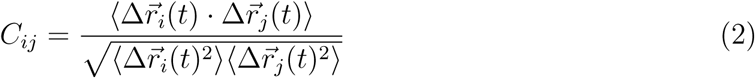

where 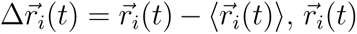 is the position vector of the *i*th residue’s *α*-carbon at time *t* and 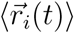 is the *i*th node’s mean position.^46^

Dynamic network analysis was performed using the dynetan 1.0.1 python package. ^45^ The network was created by assigning a node to the *α*-carbon of each residue and connecting nodes with edges if two nodes were within 4.5 Å for over 75% of equilibrated frames. After removing edges between nearest neighbours, the remaining edges were weighted by their normalized correlation between nodes:

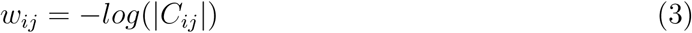

where *w_ij_* is the weight of an edge between nodes *i* and *j* and *C_ij_* is the normalized correlation. *w_ij_* represents the probability of information transfer across an edge. Inclusion of nearest neighbours did not significantly affect assigned communities or optimal paths.

The shortest distance between all nodes was calculated using the Floyd-Warshall algorithm, as implemented in the dynetan package. Communities were clustered using Louvain heuristics until a modularity metric was maximized. The modularity measures the probability difference between intra- and inter-community edges. Modularity values of all systems were about 0.75, which is slightly higher than scores in typical real-world networks (0.4-0.7).^80^

## Author Contributions

E.L.: conceptualization, methodology, investigation, visualization, writing - original draft. S.R.: conceptualization, methodology, supervision, writing - review & editing.

## Data Availability Statement

MD simulation data (initial coordinates and simulation trajectories) are provided for all systems as a Zenodo repository accessible here: https://zenodo.org/records/13730633. Data underlying figures are also available in this Zenodo repository.

## Acknowledgement

The authors thank Dr. Natalia Díaz from the University of Oviedo for providing the structure of the SARS-CoV-2 M^pro^ with the pentapeptide substrate mimic. This research was supported by a Connaught New Researcher Award to S.R. and a Natural Science and Engineering Research Council of Canada (NSERC) Discovery Grant. This research was enabled in part by support provided by Calcul Québec and the Digital Research Alliance of Canada (alliance.can.ca). Computations were performed on the Niagara supercomputer at the SciNet HPC Consortium. SciNet is funded by: the Canada Foundation for Innovation; the Government of Ontario; Ontario Research Fund – Research Excellence; and the University of Toronto.

## Supporting Information Available

Supporting Information: Additional simulation methods, including system setup and analysis; B-factors and crystal contacts; comparison of crystal structures; RMSF profiles; sequence alignment; dimensionality reduction and clustering analysis; dynamic network analysis; comparison of monomer and dimer ensembles; analysis of SARS-CoV and MERS-CoV M^pro^; principal component analysis; analysis of loop states in all systems; descriptions of Supporting Videos; References.

