## Supporting Information for "The Conformational Space of the SARS-CoV-2 Main Protease Active Site Loops is Determined by Ligand Binding and Interprotomer Allostery"

### Contents

**1 Supporting Text** . . . . . . . . . . . . . . . . . . . . . . . . . . . . . . **S-5**

1.1 Creating the Substrate-$M^{pro}$ Complex . . . . . . . . . . . . . . . . . . . . S-5

1.2 Delineation of the Equilibration Period and Sampling . . . . . . . . . . . . . . S-5

1.3 Structural Analysis . . . . . . . . . . . . . . . . . . . . . . . . . . . . S-6

**2 Supporting Figures** . . . . . . . . . . . . . . . . . . . . . . . . . . . . **S-8**

**3 Supporting Tables** . . . . . . . . . . . . . . . . . . . . . . . . . . . . **S-22**

**4 Supporting Videos** . . . . . . . . . . . . . . . . . . . . . . . . . . . . **S-30**

**References** . . . . . . . . . . . . . . . . . . . . . . . . . . . . . . . . **S-30**

### List of Figures

S1 B-factors and crystal contacts . . . . . . . . . . . . . . . . . . . . . . . . S-8

S2 Comparison of 6 SARS-CoV-2 $M^{pro}$ apo crystal structures . . . . . . . . . . S-9

S3 Free energy plotted against the distance between the upper or lower loop tip ($Leu_{50}$ or $Arg_{188}$ respectively) and $Pro_9$ . . . . . . . . . . . . . . . . . . . . S-10

S4 RMSF of catalytic dyad atoms in different upper loop states from the apo dimer simulations . . . . . . . . . . . . . . . . . . . . . . . . . . . . S-11

S5 $Arg_{188}$-$Pro_9$ distance and $Arg_{188}$ $\psi$ angle time series from the apo dimer simulations . . . . . . . . . . . . . . . . . . . . . . . . . . . . S-12

S6 Sequence alignment of SARS-CoV variants . . . . . . . . . . . . . . . . . . S-13

S7 Characterization of the apo and bound protomers of the one-substrate system through identification of active site loop states . . . . . . . . . . . . . . S-14

S8 Paths connecting the substrate and the active site of the apo protomer in the one-substrate system . . . . . . . . . . . . . . . . . . . . . . . . . . S-15

S9 UMAP projections and HDBSCAN clusters of the upper and lower loops in the SARS-CoV-2 $M^{pro}$ monomer, SARS-CoV $M^{pro}$ dimer, and MERS-CoV $M^{pro}$ dimer . . . S-16
S10 Domain III positional shift between the SARS-CoV-2 $M^{pro}$ monomer and dimer S-17
S11 Dynamic network analysis of SARS-CoV and MERS-CoV $M^{pro}$ . . . S-18
S12 Difference in the hydrophobic burial of residues 285 and 286 in SARS-CoV and SARS-CoV-2 . . . S-19
S13 Assessment of the sampling quality of the apo dimer simulations . . . S-20
S14 Principal component analysis of upper and lower loops in apo dimer simulations S-21

### List of Tables

S1 Dependence of loop state frequencies on the state of the opposite loop . . . S-22
S2 Top 10 edges of highest betweenness between protomers in the SARS-CoV-2 one-substrate system dynamic network analysis . . . S-23
S3 Residues with the most connections between protomers in the SARS-CoV-2 one-substrate system as shown through a dynamic network analysis . . . S-23
S4 Average distance from either the upper or lower loop tip (C$\alpha$ of $Leu_{50}$ or $Arg_{188}$) to the C$\alpha$ of $Pro_9$ in open, intermediate, and closed states for the SARS-CoV-2 $M^{pro}$ systems . . . S-24
S5 Average distance from either the upper or lower loop tip ($Leu_{50}$ or $Arg_{188}$) to $Pro_9$ in all UMAP and HDBSCAN clusters for the SARS-CoV-2 $M^{pro}$ apo dimer and 2-substrate systems . . . S-25
S6 Average distance from either the upper or lower loop tip ($Leu_{50}$ or $Arg_{188}$) to $Pro_9$ in all UMAP and HDBSCAN clusters for the SARS-CoV-2 $M^{pro}$ 1-substrate system . . . S-26

S7 Average distance from either the upper or lower loop tip to $Pro_9$ in open, intermediate, and closed states for the SARS-CoV-2 $M^{pro}$ monomer and SARS-CoV and MERS-CoV systems . . . . . . . . . . . . . . . . . . . . . . . . . . . . S-27
S8 Average distance from either the upper or lower loop tip to $Pro_9$ in all UMAP and HDBSCAN clusters for the SARS-CoV and MERS-CoV systems . . . . S-28
S9 Average distance from the upper or lower loop tip ($Leu_{50}$ or $Arg_{188}$) to $Pro_9$ in all UMAP and HDBSCAN clusters for the SARS-CoV-2 apo monomer . . S-29

### 1 Supporting Text

#### 1.1 Creating the Substrate-$M^{pro}$ Complex

The structure used for the substrate-bound $M^{pro}$ simulations was obtained from Dimas Suárez and Natalia Díaz (Universidad de Oviedo), who built the structure based on the crystal structure PDB 6LU7 as described in Suárez and Díaz (2020).[S1] The procedure they used is summarized in brief here. The crystal structure (PDB 6LU7) contains the SARS-CoV-2 $M^{pro}$ dimer bound to a peptidomimetic N3 inhibitor.[S2] $M^{pro}$ cleaves polyproteins at various peptide sequences with a conserved glutamine at the P1 site, where cleavage occurs at the peptide bond between P1-P1'.[S3,S4] The cleavage site between $M^{pro}$ and nonstructural protein 4 in the polyprotein corresponds to a -P4-P3-P2-P1–P1'- sequence of -Ala-Val-Leu-Gln–Ser-, which resembles the N3 inhibitor. The inhibitor included an -Ala-Val-Leu- portion in the -P4-P3-P2- sites that was left unchanged. A segment resembling a Gln was located in the P1 site, which was modified into a Gln. The rest of the inhibitor was removed, and a Ser residue was built in its place along the original inhibitor backbone at the P1' site. The bond between the catalytic $Cys_{145}$ and the inhibitor was removed, and N-terminal acetyl- and C-terminal N-methyl amide capping groups were built into the termini. Histidine residues were protonated to be consistent with a pH of 7.0. For a full description of the procedure to build the substrate, refer to the study by Suárez and Díaz.[S1]

#### 1.2 Delineation of the Equilibration Period and Sampling

Trajectories were evaluated based on four metrics of equilibration: RMSD, radius of gyration ($R_{gyr}$), number of hydrogen bonds, and solvent accessible surface area (SASA). These results are shown in Figure 1D for the apo dimer. Trajectories were determined to have reached equilibrium after 0.5 µs of sampling based on these metrics. The next 1 µs of simulation was used for analysis of the equilibrium ensembles. Simulations of the apo monomer were determined to have reached equilibrium after 2 µs. The next 1 µs was then analyzed. Only portions of the one- and two-substrate bound systems where the substrate had not dissociated were used for analysis.

Sampling was assessed by splitting the equilibrated portions of trajectories into two halves and comparing observables to determine if their mean values were consistent within error. The structural metrics we used for this assessment were $R_{gyr}$, number of hydrogen bonds, and solvent accessible surface area (SASA). This assessment is shown for the apo dimer system as a representative example in Figure S13. As an additional approach to assess the extent of conformational sampling in the simulations, we used principal component analysis (PCA). This analysis was carried out for the upper and lower loops of each trajectory of the apo dimer. The results show that conformations of the loops are revisited during the trajectories, which is another indication of good sampling (Figure S14), in addition to the comparison of the global metrics shown in Figure S13.

#### 1.3 Structural Analysis

All analysis of the trajectories was carried out using a combination of GROMACS 2019.1[S5] and the MDAnalysis 1.0.0 Python package.[S6,S7] Hydrogen bonds were identified using the gmx hbond module in GROMACS and were determined based on a 30° cut-off for the hydrogen-donor-acceptor angle and a 0.35 nm cut-off for the donor-acceptor distance. OH and NH groups were considered donors while O and N were considered acceptors. A water bridge was defined as present if hydrogen bond donors/acceptors of all potential bridging residues were within 3.5 Å of the same water oxygen. Visual Molecular Dynamics (VMD) 1.9.4[S8] and UCSF ChimeraX 1.7.1[S9] were used for the visualization of structures and simulation trajectories. Jensen-Shannon distances between ensembles were calculated using PENSA[S10] using either dihedral angles or $\alpha$-carbon distances as features. In the two-substrate system, analysis was run on equilibrated sections of all trajectories where both substrates were bound to the active site. Standard errors of the mean and standard deviations were calculated using the scipy 1.5.4 Python package.[S11] UMAP, and HDBSCAN were performed using the scikit-learn 0.23.2,[S12] umap-learn 0.4.6,[S13] and hdbscan 0.8.26[S14] Python packages respectively.

Free energy profiles of the different loop states shown in Figure S3 were calculated using the formula $G(x) = -k_B T \log p(x)$,[S15] where $p(x)$ is the distribution of the distance between the loop tip and $\text{Pro}_9$ ($\text{Leu}_{50}$-$\text{Pro}_9$ distance in the upper loop and $\text{Arg}_{188}$-$\text{Pro}_9$ distance in the lower loop). $p(x)$ was computed using a histogram estimator with 100 bins. The standard error in $p(x)$ was computed using bootstrapping with n = 1000.

#### 2 Supporting Figures

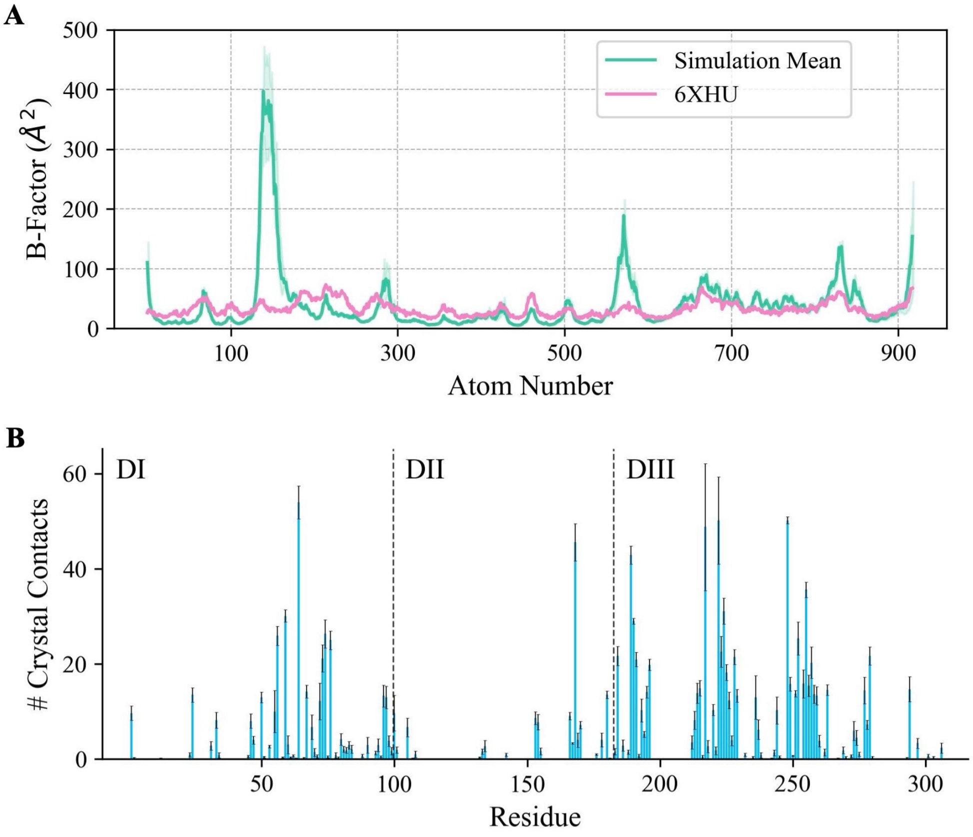


Figure S1: **B-factors and crystal contacts.** (A) B-factor comparison between the SARS-CoV-2 M$^{pro}$ apo dimer simulations (green) and crystal structure (pink). Green shading indicates standard error. The upper and lower loops comprise atom numbers 129-161 and 549-581, respectively. (B) Crystal contacts from 6 apo SARS-CoV-2 M$^{pro}$ structures (6WQF, 6XHU, 6M03, 6YB7, 6Y2E, 6WTM). Crystal contacts are defined as any contacts within 5 Å between heavy atoms from separate unit cells in a crystal structure. The mean value of the number of contacts is shown for each residue and the error bars indicate the standard error of the mean.

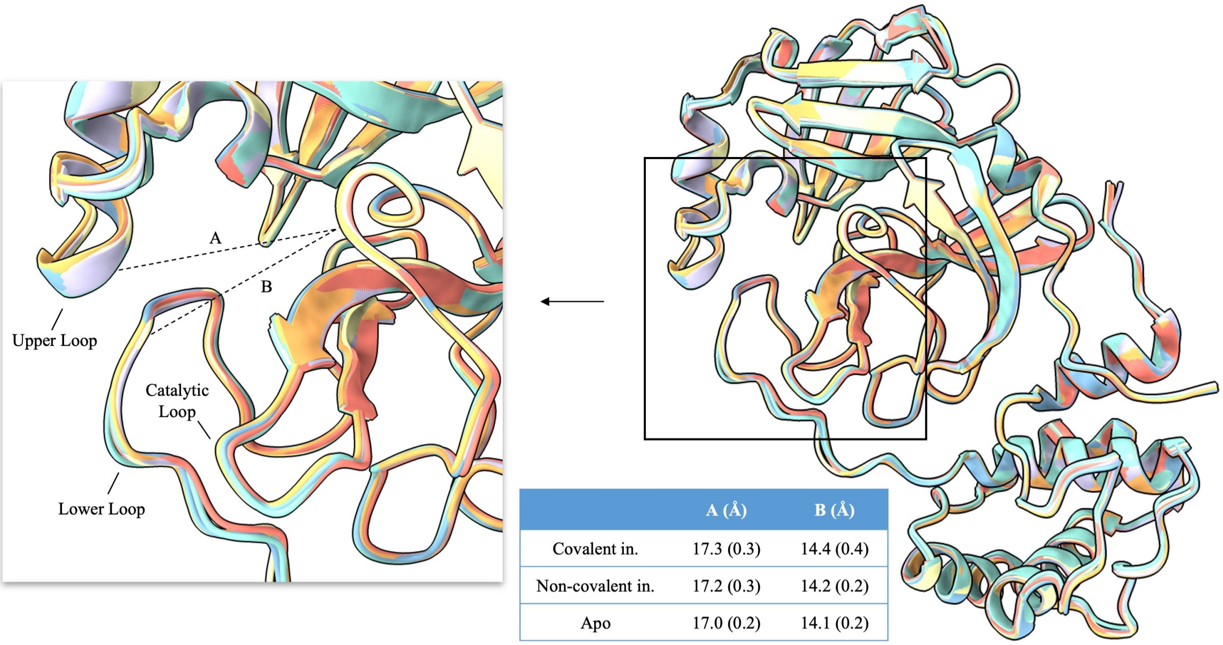


| | A (Å) | B (Å) |
|---|---|---|
| Covalent in. | 17.3 (0.3) | 14.4 (0.4) |
| Non-covalent in. | 17.2 (0.3) | 14.2 (0.2) |
| Apo | 17.0 (0.2) | 14.1 (0.2) |

Figure S2: **Comparison of 6 SARS-CoV-2 $M^{pro}$ apo crystal structures.** Six structures of an apo monomer are shown (6YB7 in pink, 6Y2E in turquoise, 6WTM in lime, 6M03 in ice blue, 6WQF in yellow, 6XHU in cyan). A close-up view of the active site is shown on the left. Dashed lines indicate distances between the $\alpha$-carbons of $Cys_{145}$ and either $Leu_{50}$ (line A) or $Gln_{189}$ (line B). The table provided in the inset provides the mean value of these distances computed from 116 crystal structures with covalent inhibitors, 96 structures with non-covalent inhibitors, and 25 apo structures of $M^{pro}$. Standard deviations of these distances are shown in brackets.

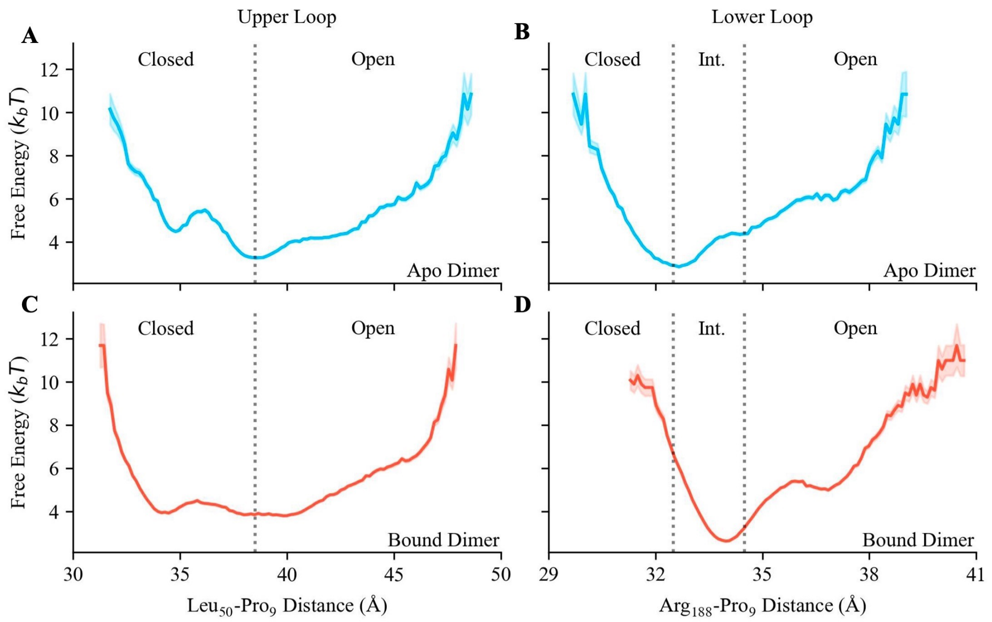


Figure S3: **Free energy plotted against the distance between the upper or lower loop tip (Leu$_{50}$ or Arg$_{188}$ respectively) and Pro$_9$.** (A) and (B) show the free energy of the upper and lower loops in different states in the apo dimer, while (C) and (D) show the free energy of the upper and lower loops in different states in the bound dimer (two-substrate system). The different states of each loop are labelled on each plot, with dotted lines indicating the distance criteria for each state used during dimensionality reduction and categorization. Shaded areas represent the standard error computed from bootstrapping with $n = 1000$. The scale of the y-axis is the same in all plots. The free energy calculation is described in SI Section 1.3.

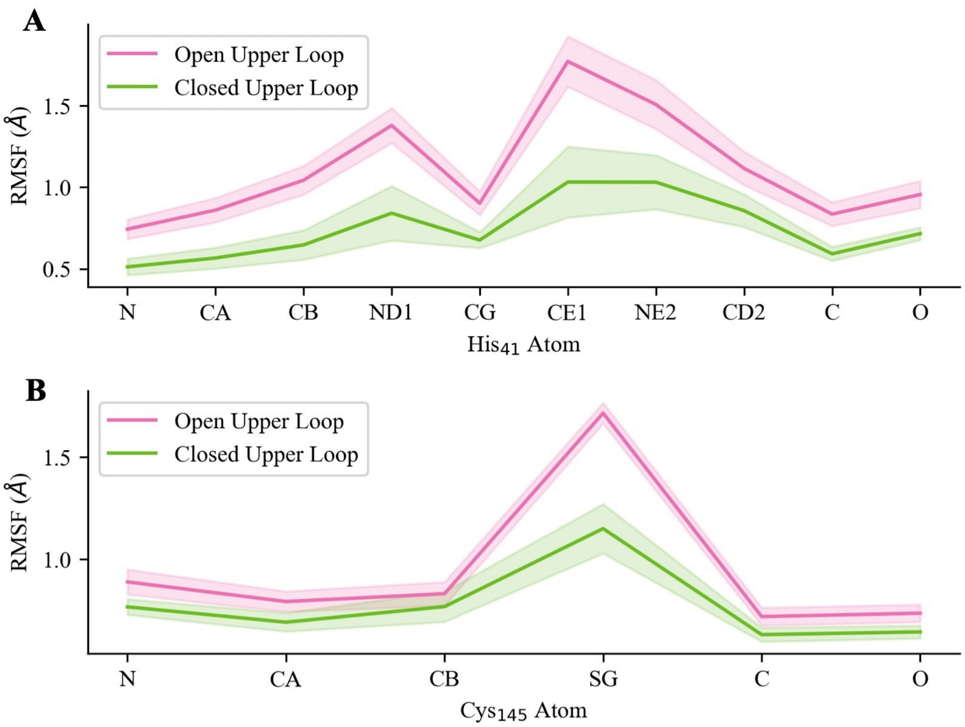


Figure S4: **RMSF of catalytic dyad atoms in different upper loop states from the apo dimer simulations.** Comparisons of RMSF per atom in (A) $His_{41}$ and (B) $Cys_{145}$ in open and closed upper loop states. Shaded areas denote standard error of the mean.

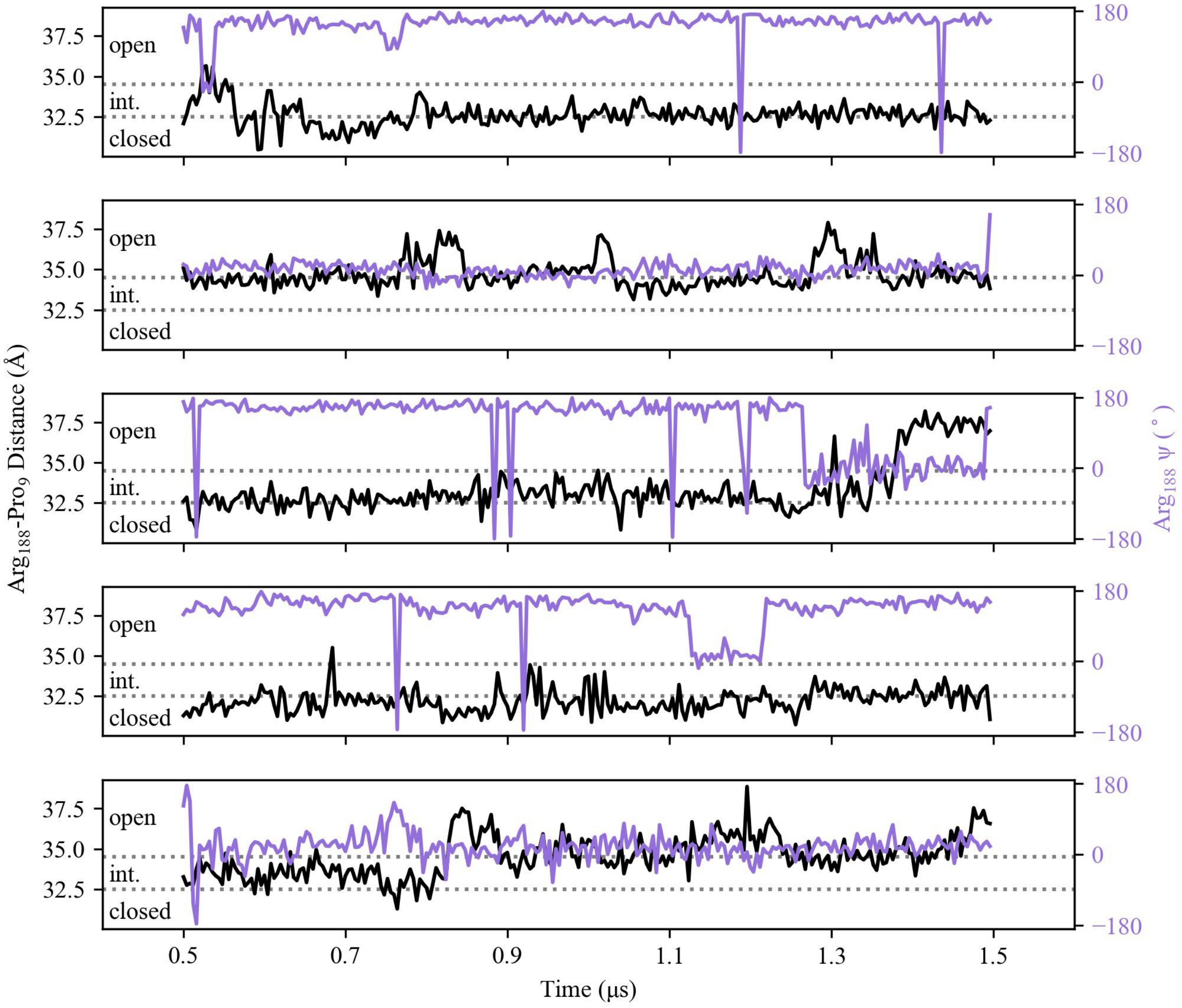


Figure S5: **$Arg_{188}$-$Pro_9$ distance and $Arg_{188}$ $\psi$ angle time series from the apo dimer simulations.** The $Arg_{188}$-$Pro_9$ distance, which was used to classify lower loop states, is plotted alongside the $Arg_{188}$ $\psi$ angle to show the trend between the orientation of the $Arg_{188}$ side chain and lower loop state. The different states of the lower loop are labelled on each plot, with dotted lines indicating the distance criteria for each state used during dimensionality reduction and categorization. Time series are shown for 5 replicates.

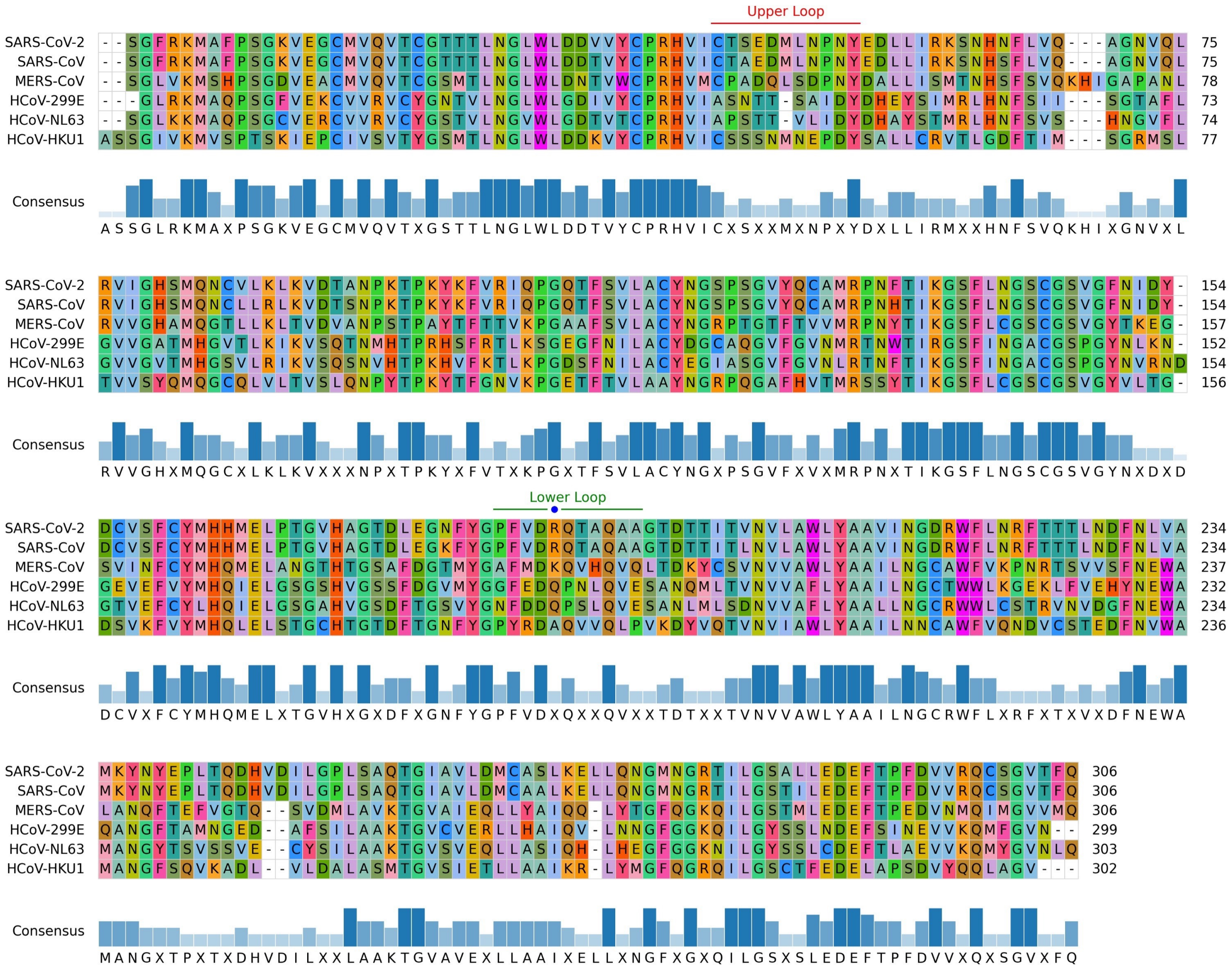

Figure S6: **Sequence alignment of SARS-CoV variants.** Alignment of HCoV-299E, HCoV-NL63, SARS-CoV-2, SARS-CoV, MERS-CoV, and HCoV-HKU1 M$^{pro}$s using Clustal Omega,[S16] visualized using pyMSAviz.[S17] Although not involved in this study, HCoV-299E, HCoV-NL63, and HCoV-HKU1 are related human coronaviruses included here to show the conservation of residues in the upper and lower loops. Boxes are coloured by residue. Bars below each residue show the highest consensus residue percentage. The upper and lower loops are indicated. The blue circle in the lower loop indicates Arg$_{188}$ in the SARS-CoV-2 M$^{pro}$. Sequence identities for each variant compared to SARS-CoV-2 are as follows: HCoV-299E (41.14%), HCoV-NL63 (44.04%), SARS-CoV (96.08%), MERS-CoV (50.83%), HCoV-HKU1 (49.33%).

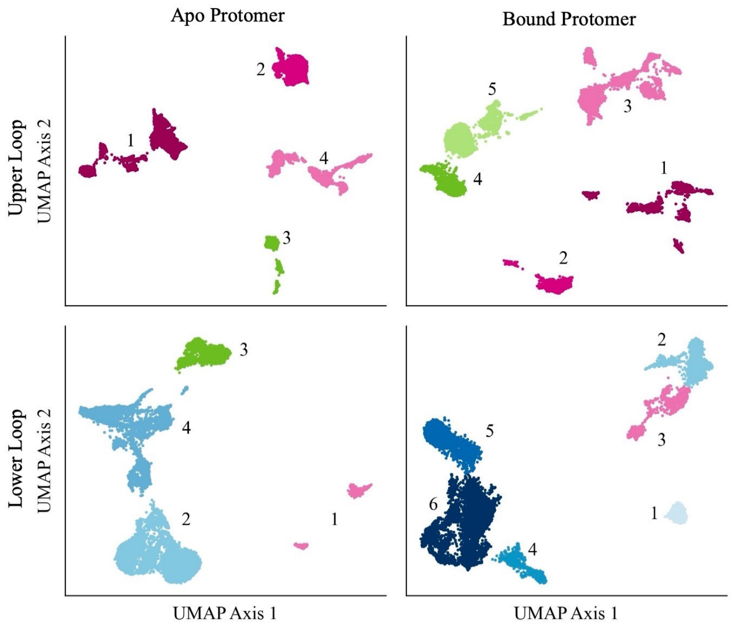


Figure S7: **Characterization of the apo and bound protomers of the one-substrate system through identification of active site loop states.** UMAP projections and HDBSCAN clusters of the SARS-CoV-2 M$^{pro}$ upper loop (upper panels) and lower loop (lower panels) in the apo protomer (left panels) and bound protomer (right panels) of the one-substrate system. Numbers within the UMAP plots signify the cluster numbers. Clusters coloured in pink, blue, and green shades denote open, intermediate, and closed loop states, respectively. The different shades of each colour are assigned arbitrarily.

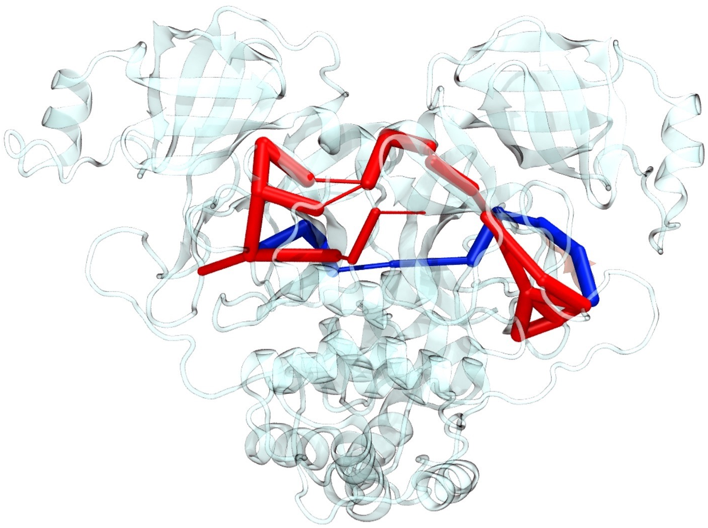

Figure S8: **Paths connecting the substrate and the active site of the apo protomer in the one-substrate system.** The top five optimal paths are shown. Edge thicknesses are weighted by their correlation score. The blue path is the most optimal path, while the red ones are the next four most optimal paths. Note that some of the red paths overlap because they include some of the same residues.

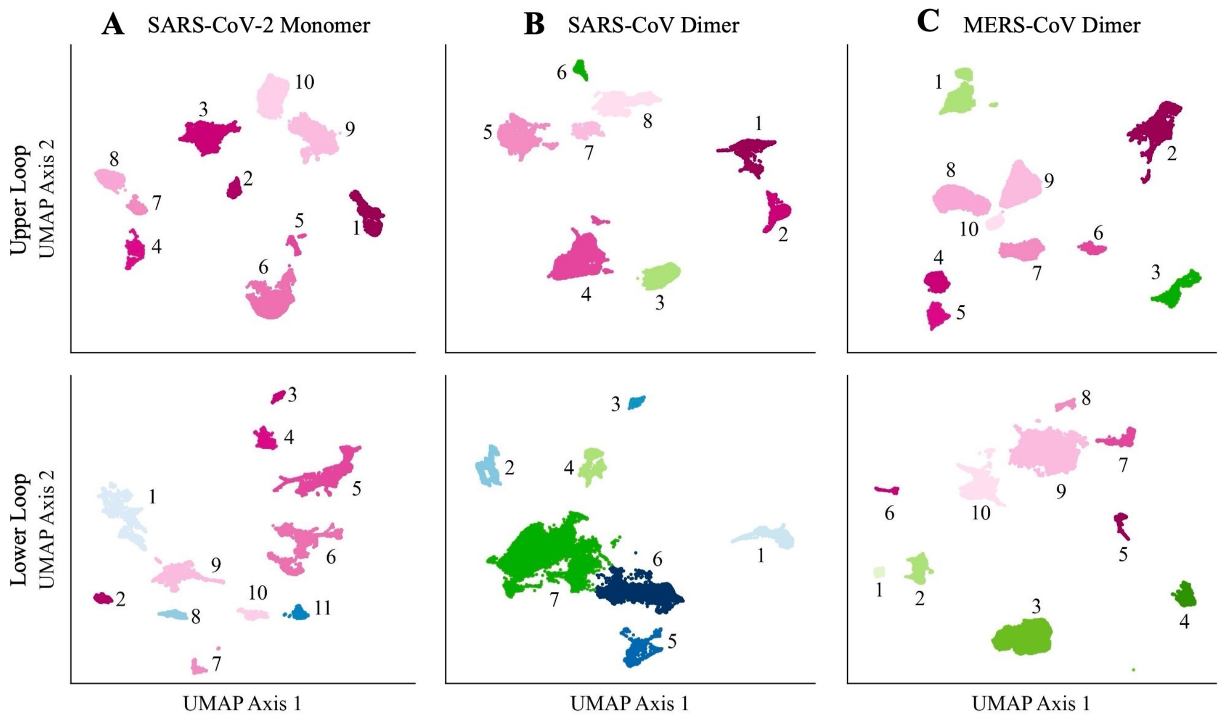


Figure S9: **UMAP projections and HDBSCAN clusters of the upper and lower loops in the SARS-CoV-2 M$^{pro}$ monomer, SARS-CoV M$^{pro}$ dimer, and MERS-CoV M$^{pro}$ dimer.** (A) SARS-CoV-2 M$^{pro}$ monomer, (B) SARS-CoV M$^{pro}$ dimer, and (C) MERS-CoV M$^{pro}$ dimer projections and clusters. UMAP representations are made using the entire equilibrated dataset of each system (five independent trajectories of 1.5 µs each, excluding the first 0.5 µs for equilibration). Clustering parameters were chosen to maximize the relative validity score. Numbers within the UMAP plots signify each cluster. Clusters coloured in pink, blue, and green shades denote open, intermediate, and closed loop states, respectively. The different shades of each colour are assigned arbitrarily. Clusters are categorized as closed or open based on the distance between the loop tip and the N-terminus, which is used as a reference point due to its stability, in combination with visual inspection. The distances used to categorize these states can be found in Table S8 and Table S9.

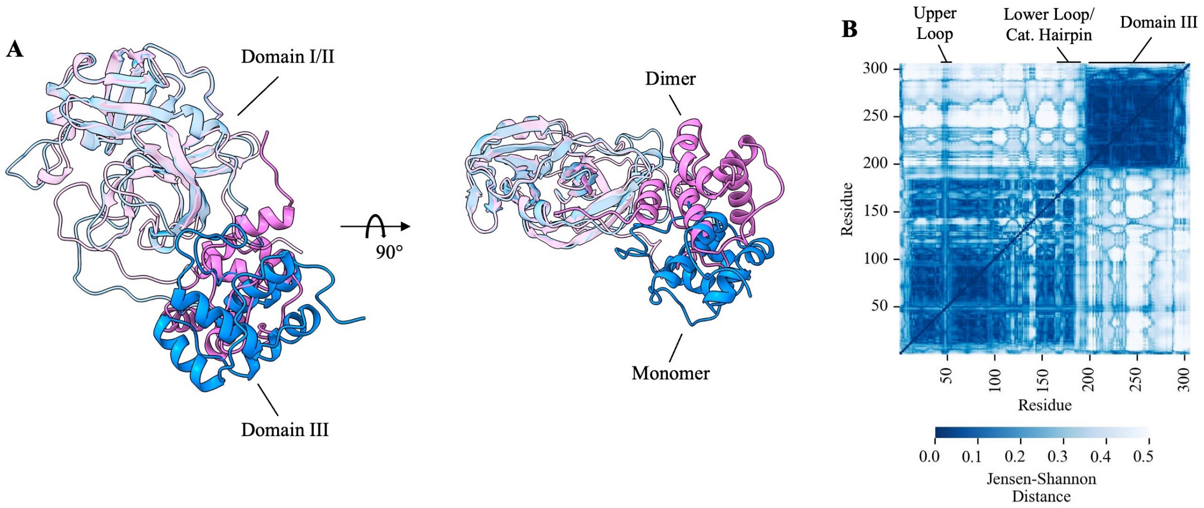


Figure S10: **Domain III positional shift between the SARS-CoV-2 M$^{pro}$ monomer and dimer.** (A) Structural views of the Domain III shift. The dimer and monomer structures, taken at random timepoints from the apo dimer and apo monomer simulations, are in pink and blue, respectively. Structures were aligned by Domains I and II. (B) Jensen-Shannon distance values between the monomer and apo dimer ensembles using inter-C$\alpha$ distances for featurization.

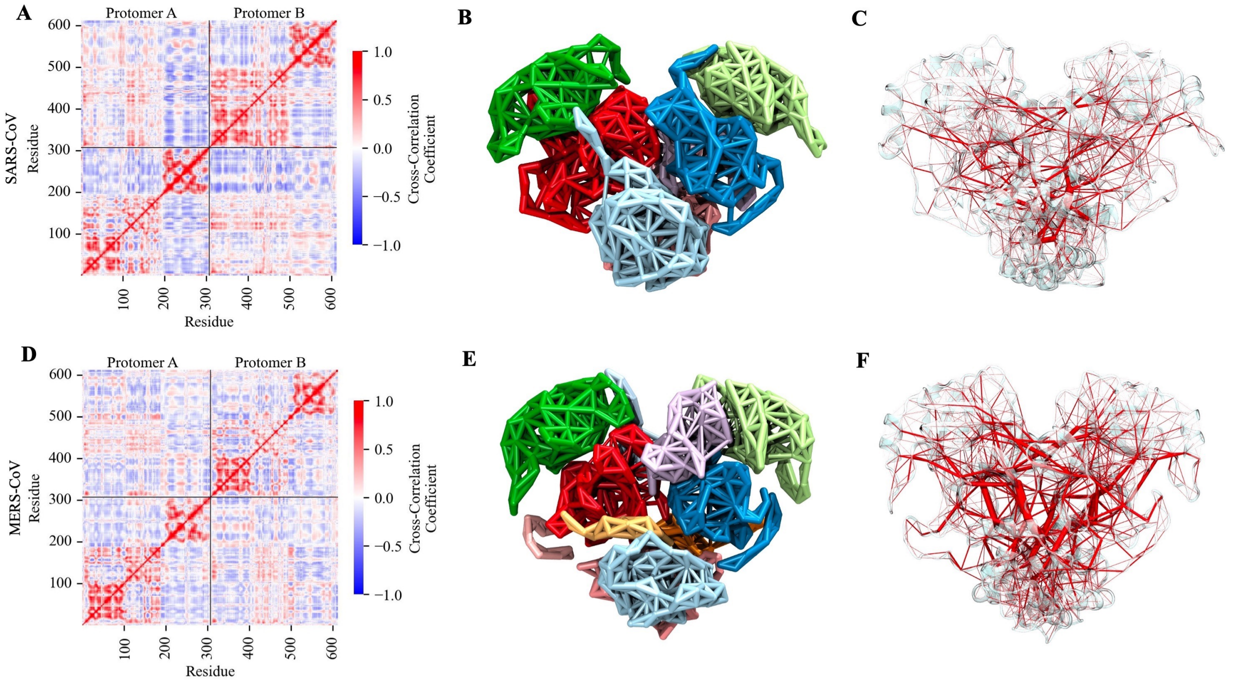


Figure S11: **Dynamic network analysis of SARS-CoV and MERS-CoV M$^{pro}$.** (A), (D) Cross-correlation coefficient matrices. Values close to 1, -1, and 0 show correlation, anti-correlation, and no correlation, respectively. (B), (E) Community breakdown of the M$^{pro}$ in both variants. Communities are indicated by different colours. (C), (F) Network representation of the M$^{pro}$ with edges weighted by betweenness. To improve visualization, the betweenness values are normalized by dividing each raw value by the highest betweenness in its respective system. As a result, edge thicknesses are not directly comparable between systems. Results for SARS-CoV and MERS-CoV M$^{pro}$ are shown in the top and bottom rows, respectively.

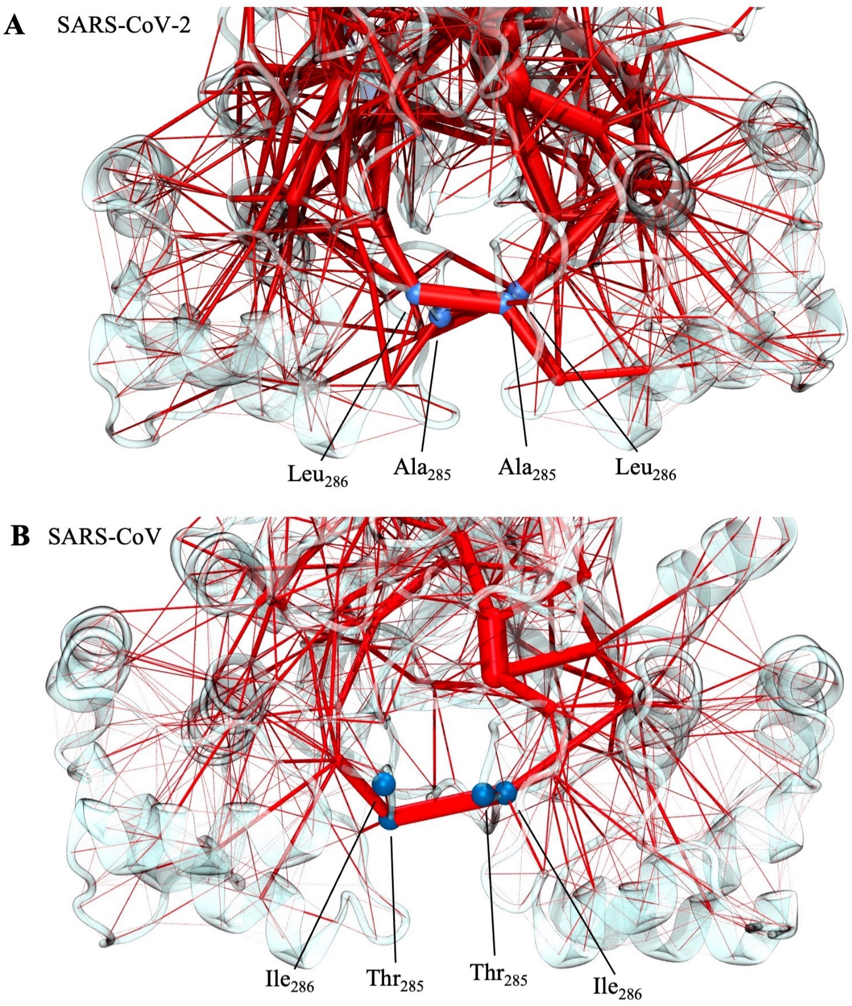


Figure S12: **Difference in the hydrophobic burial of residues 285 and 286 in SARS-CoV and SARS-CoV-2.** Betweenness representations from the dynamic network analyses of (A) SARS-CoV $M^{pro}$ and (B) SARS-CoV-2 $M^{pro}$, zoomed in on the dimerization interface between Domain III's. Red edges show the betweenness between two residues, with thicker lines indicating higher betweenness (stronger potential for communication). In (A) SARS-CoV, residues $Thr_{285}$ and $Ile_{286}$ ($C\alpha$ atoms shown in blue spheres) form a loose bridge between protomers. In (B) SARS-CoV-2, equivalent residues $Ala_{285}$ and $Leu_{286}$ are buried closer together due to hydrophobic interactions, resulting in more edges of high betweenness between the two protomers not seen in SARS-CoV.

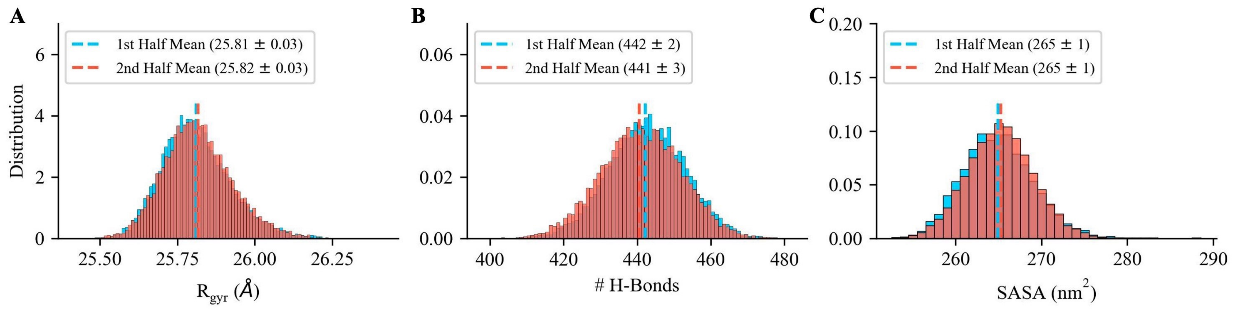


Figure S13: **Assessment of the sampling quality of the apo dimer simulations.** The equilibrated portions of the trajectories were split into the first and second halves. The $R_{gyr}$ (A), number of hydrogen bonds (B), and SASA (C) in the first and second halves were compared to evaluate sampling. For all of these structural properties, the mean values computed based on the first and second half of the simulation are consistent within error.

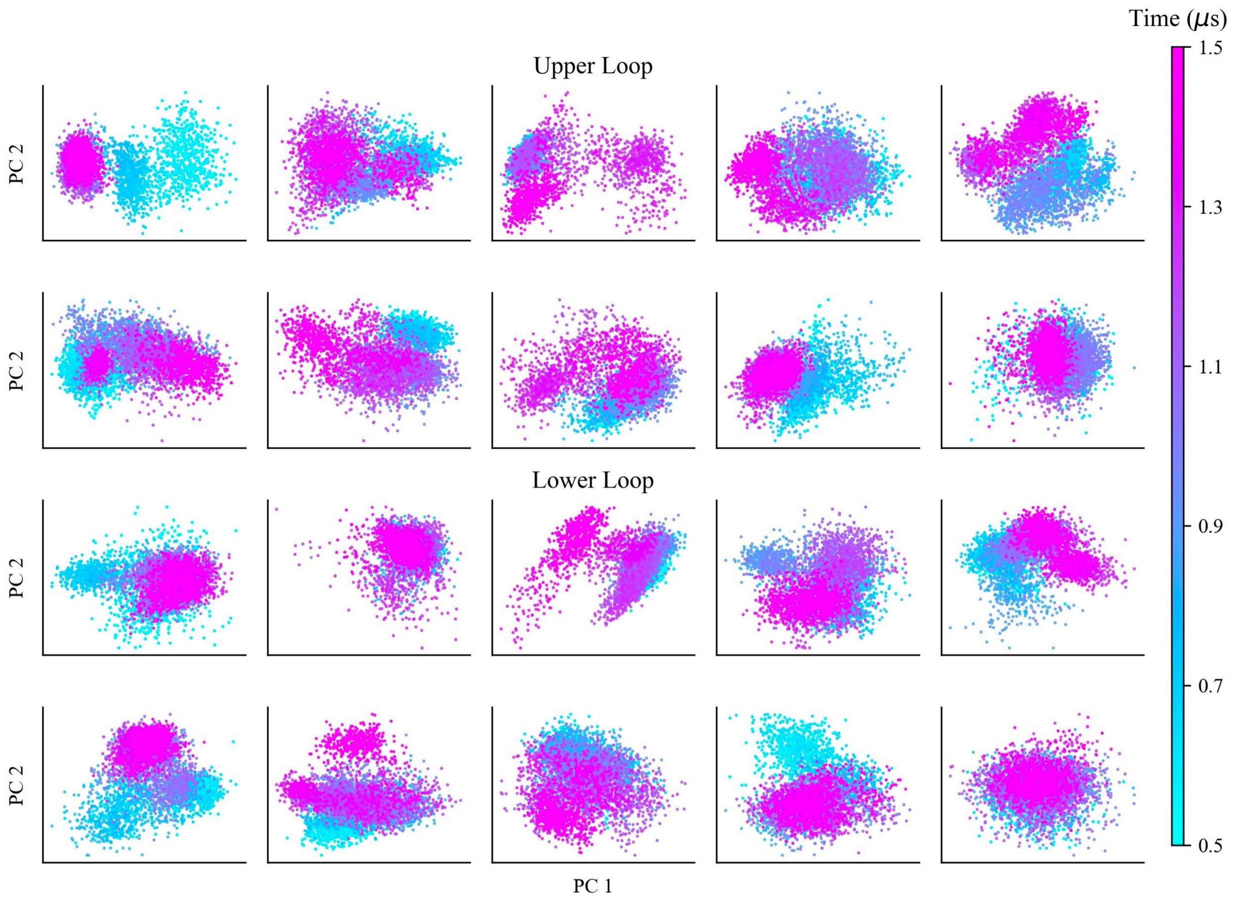


Figure S14: **Principal component analysis of upper and lower loops in apo dimer simulations.** Projections of the first two principal components of all upper (residues 44-54) and lower loop (residues 184-194) heavy atom positions from five SARS-CoV-2 M$^{pro}$ apo dimer simulations, after alignment excluding the active site loops and termini. Principal component analysis was carried out for each trajectory separately. Each dimer contains two upper and lower loops, resulting in ten trajectories in total. Frames are coloured by their time in the trajectory, with the first 0.5 μs excluded for equilibration.

### 3 Supporting Tables

Table S1: **Dependence of loop state frequencies on the state of the opposite loop.** Occupancy percentages were computed from UMAP and HDBSCAN analysis of the SARS-CoV-2 apo $M^{pro}$ dimer simulations. The table shows the frequency (in %) of each possible loop state given the opposite loop's state. The difference in the frequency of that loop's state from the average (ie. combined loop state percentages across the whole trajectory from Figure 2) is also provided in the third column of the table.

| Lower loop state if upper loop is open | | |
|---|---|---|
| Lower Loop State | Occupancy (%) | Difference from Average (%) |
| Open | 32.0 | +4.5 |
| Intermediate | 38.3 | -2.5 |
| Closed | 29.7 | -1.9 |

| Lower loop state if upper loop is closed | | |
|---|---|---|
| Lower Loop State | Occupancy (%) | Difference from Average (%) |
| Open | 10.1 | -17.4 |
| Intermediate | 48.2 | +7.4 |
| Closed | 41.7 | +10.1 |

| Upper loop state if lower loop is open | | |
|---|---|---|
| Upper Loop State | Occupancy (%) | Difference from Average (%) |
| Open | 93.6 | +14.5 |
| Closed | 6.4 | -14.5 |

| Upper loop state if lower loop is intermediate | | |
|---|---|---|
| Upper Loop State | Occupancy (%) | Difference from Average (%) |
| Open | 78.5 | +0.6 |
| Closed | 21.5 | -0.6 |

| Upper loop state if lower loop is closed | | |
|---|---|---|
| Upper Loop State | Occupancy (%) | Difference from Average (%) |
| Open | 76.6 | -2.5 |
| Closed | 23.4 | +2.5 |

Table S2: **Top 10 edges of highest betweenness between protomers in the SARS-CoV-2 one-substrate system dynamic network analysis.** The higher the betweenness, the stronger the communication between residues. Several N-terminal residues (residues 1-9, shown in bold) populate the list, consistent with the idea that the active site is shaped in part by the opposite protomer's N-terminus and signifying the importance of the N-terminus in interprotomer communication. Correlation coefficients for each pair of residues is also shown, indicating the degree of correlated movement between the two residues throughout the simulations (with higher values signifying stronger correlation).

| Bound Protomer Residue | Apo Protomer Residue | Betweenness | Correlation Coefficient |
|---|---|---|---|
| **$Arg_4$** | $Ser_{139}$ | 0.047 | 0.22 |
| $Glu_{14}$ | $Lys_{12}$ | 0.042 | 0.22 |
| $Lys_{137}$ | **$Arg_4$** | 0.025 | 0.27 |
| $Tyr_{126}$ | **$Lys_5$** | 0.020 | 0.39 |
| $Gly_{138}$ | **$Phe_3$** | 0.019 | 0.24 |
| $Glu_{14}$ | **$Pro_9$** | 0.019 | 0.14 |
| **$Gly_2$** | $Leu_{141}$ | 0.018 | 0.22 |
| $Phe_{140}$ | $Gln_{299}$ | 0.018 | 0.36 |
| $Ala_{285}$ | $Leu_{286}$ | 0.018 | 0.45 |
| $Leu_{141}$ | $Arg_{298}$ | 0.018 | 0.40 |

Table S3: **Residues with the most connections between protomers in the SARS-CoV-2 one-substrate system as shown through a dynamic network analysis.** A = apo protomer, B = bound protomer.

| **Residue** | **# Connections** |
|---|---|
| $Met_{6A}$ | 7 |
| $Ser_{139B}$, $Pro_{9A}$ | 6 |
| $Val_{125B}$, $Arg_{4A}$, $Ser_{1A}$ | 5 |
| $Met_{6B}$, $Pro_{9B}$, $Gly_{138B}$ | 4 |
| $Ser_{1B}$, $Ala_{7B}$, $Ser_{10B}$, $Glu_{14B}$, $Gly_{124B}$, $Gln_{127B}$, $Leu_{141B}$, $Leu_{286B}$, $Phe_{305B}$, $Val_{125A}$, $Ser_{123A}$, $Gly_{11A}$, $Gln_{299A}$ | 3 |

Table S4: **Average distance from either the upper or lower loop tip (C$\alpha$ of Leu$_{50}$ or Arg$_{188}$) to the C$\alpha$ of Pro$_9$ in open, intermediate, and closed states for the SARS-CoV-2 M$^{pro}$ systems.** Distances provided are averages over all conformations in clusters categorized as open/intermediate/closed. These values were used in combination with visual inspection to categorize UMAP and HDBSCAN clusters. Pro$_9$ was chosen as a reference point because it exhibits minimal movement in the trajectories. Standard deviation is provided in brackets for each distance. The Leu$_{50}$-Pro$_9$ and Arg$_{188}$-Pro$_9$ distances from six crystal structures (6YB7, 6Y2E, 6WTM, 6M03, 6WQF, and 6XHU) were used as a rough guideline for the classification of states. The mean Leu$_{50}$-Pro$_9$ and Arg$_{188}$-Pro$_9$ distances were 37.14 $\pm$ 0.03 and 32.73 $\pm$ 0.02 Å respectively. The upper and lower loops are in closed and intermediate states respectively for all crystal structures.

| | **Apo Dimer** | **2 Sub. Dimer** | **1 Sub. Dimer Bound Protomer** | **1 Sub. Dimer Apo Protomer** |
|---|---|---|---|---|
| **Upper Loop State** | **Mean Pro$_9$-Leu$_{50}$ Distance (Å)** | | | |
| **Open** | 40.1 (2.4) | 39.8 (2.4) | 39.4 (2.6) | 42.5 (2.7) |
| **Closed** | 37.1 (1.8) | 36.0 (2.1) | 34.0 (0.9) | 36.0 (3.1) |
| **Lower Loop State** | **Mean Pro$_9$-Arg$_{188}$ Distance (Å)** | | | |
| **Open** | 34.7 (1.3) | 36.4 (1.1) | 35.7 (0.9) | 35.5 (1.2) |
| **Intermediate** | 32.7 (0.6) | 34.0 (0.6) | 34.2 (0.5) | 32.8 (0.4) |
| **Closed** | 32.1 (0.6) | - | - | 32.3 (0.7) |

Table S5: **Average distance from either the upper or lower loop tip ($Leu_{50}$ or $Arg_{188}$) to $Pro_9$ in all UMAP and HDBSCAN clusters for the SARS-CoV-2 $M^{pro}$ apo dimer and 2-substrate systems.** These values were used in combination with visual inspection to categorize UMAP and HDBSCAN clusters. $Pro_9$ was chosen as a reference point because it exhibits minimal movement in the trajectories. Standard deviations are provided in brackets.

| SARS-CoV-2 Apo Dimer | | | SARS-CoV-2 2 Sub. Dimer | | |
|---|---|---|---|---|---|
| **Cluster** | **Upper Loop Mean $Pro_9$-$Leu_{50}$ Distance (Å)** | **State** | **Cluster** | **Upper Loop Mean $Pro_9$-$Leu_{50}$ Distance (Å)** | **State** |
| 1 | 38.2 (0.5) | Closed | 1 | 36.0 (2.4) | Closed |
| 2 | 41.3 (1.4) | Open | 2 | 39.8 (2.1) | Open |
| 3 | 37.7 (1.2) | Closed | | | |
| 4 | 38.3 (1.4) | Closed | | | |
| 5 | 39.2 (0.9) | Open | | | |
| 6 | 42.1 (2.1) | Open | | | |
| 7 | 40.3 (2.0) | Open | | | |
| 8 | 42.9 (1.4) | Open | | | |
| 9 | 38.5 (0.7) | Open | | | |
| 10 | 37.8 (0.6) | Closed | | | |
| 11 | 38.8 (1.3) | Open | | | |
| 12 | 38.9 (2.9) | Open | | | |
| 13 | 35.5 (1.7) | Closed | | | |

| **Cluster** | **Lower Loop Mean $Pro_9$-$Arg_{188}$ Distance (Å)** | **State** | **Cluster** | **Lower Loop Mean $Pro_9$-$Arg_{188}$ Distance (Å)** | **State** |
|---|---|---|---|---|---|
| 1 | 34.7 (1.5) | Open | 1 | 36.4 (1.1) | Open |
| 2 | 32.2 (0.5) | Closed | 2 | 34.2 (0.7) | Intermediate |
| 3 | 32.0 (0.6) | Closed | 3 | 34.0 (0.5) | Intermediate |
| 4 | 34.8 (0.9) | Open | | | |
| 5 | 33.1 (0.5) | Intermediate | | | |
| 6 | 32.8 (0.5) | Intermediate | | | |
| 7 | 32.2 (0.6) | Closed | | | |
| 8 | 31.6 (0.5) | Closed | | | |
| 9 | 32.4 (0.5) | Closed | | | |
| 10 | 32.2 (0.5) | Closed | | | |
| 11 | 32.7 (0.5) | Intermediate | | | |
| 12 | 32.6 (0.7) | Intermediate | | | |

Table S6: **Average distance from either the upper or lower loop tip ($Leu_{50}$ or $Arg_{188}$) to $Pro_9$ in all UMAP and HDBSCAN clusters for the SARS-CoV-2 $M^{pro}$ 1-substrate system.** These values were used in combination with visual inspection to categorize UMAP and HDBSCAN clusters. $Pro_9$ was chosen as a reference point because it exhibits minimal movement in the trajectories. Standard deviations are provided in brackets.

| | SARS-CoV-2 1 Sub. Dimer Bound Protomer | | | SARS-CoV-2 1 Sub. Dimer Apo Protomer | |
|---|---|---|---|---|---|
| **Cluster** | **Upper Loop Mean $Pro_9$-$Leu_{50}$ Distance (Å)** | **State** | **Cluster** | **Upper Loop Mean $Pro_9$-$Leu_{50}$ Distance (Å)** | **State** |
| 1 | 39.8 (1.7) | Open | 1 | 44.2 (1.9) | Open |
| 2 | 39.5 (2.1) | Open | 2 | 42.3 (1.4) | Open |
| 3 | 39.2 (3.2) | Open | 3 | 36.0 (3.1) | Closed |
| 4 | 33.9 (1.1) | Closed | 4 | 39.9 (2.5) | Open |
| 5 | 34.1 (0.9) | Closed | | | |

| Cluster | Lower Loop Mean $Pro_9$-$Arg_{188}$ Distance (Å) | State | Cluster | Lower Loop Mean $Pro_9$-$Arg_{188}$ Distance (Å) | State |
|---|---|---|---|---|---|
| 1 | 34.4 (0.5) | Intermediate | 1 | 35.5 (1.2) | Open |
| 2 | 35.3 (0.7) | Open | 2 | 32.8 (0.4) | Intermediate |
| 3 | 36.3 (0.9) | Open | 3 | 32.4 (0.6) | Closed |
| 4 | 34.2 (0.4) | Intermediate | 4 | 32.2 (0.7) | Closed |
| 5 | 34.3 (0.4) | Intermediate | | | |
| 6 | 34.0 (0.8) | Intermediate | | | |

Table S7: **Average distance from either the upper or lower loop tip to $Pro_9$ in open, intermediate, and closed states for the SARS-CoV-2 $M^{pro}$ monomer and SARS-CoV and MERS-CoV systems.** $Leu_{50}$ was used as the tip residue in the upper loop. $Arg_{188}$ was used as the tip residue in the lower loop in SARS-CoV-2 and SARS-CoV, while $Lys_{191}$ was used in MERS-CoV based on a sequence alignment (Figure S4) These values were used in combination with visual inspection to categorize UMAP and HDBSCAN clusters. $Pro_9$ was chosen as a reference point due to its minimal movement. Standard deviations are denoted in brackets.

| | **SARS-CoV-2 Apo Monomer** | **SARS-CoV Apo Dimer** | **MERS-CoV Apo Dimer** |
|---|---|---|---|
| **Upper Loop State** | **Mean $Pro_9$-$Leu_{50}$ Distance (Å)** | | |
| **Open** | 41.4 (2.8) | 40.4 (2.2) | 40.1 (1.7) |
| **Closed** | - | 37.5 (0.9) | 37.1 (1.8) |
| **Lower Loop State** | **Mean $Pro_9$-$Arg_{188}$ Distance (Å)** | | |
| **Open** | 36.8 (2.2) | - | 36.6 (2.6) |
| **Intermediate** | 34.1 (2.2) | 32.9 (0.8) | - |
| **Closed** | - | 32.1 (0.7) | 31.0 (0.9) |

Table S8: **Average distance from either the upper or lower loop tip to $Pro_9$ in all UMAP and HDBSCAN clusters for the SARS-CoV and MERS-CoV systems.** $Leu_{50}$ was used as the tip residue in the upper loop. $Arg_{188}$ was used as the tip residue in the lower loop in SARS-CoV, while $Lys_{191}$ was used in MERS-CoV based on a sequence alignment (Figure S4). These values were used in combination with visual inspection to categorize UMAP and HDBSCAN clusters. $Pro_9$ was chosen as a reference point due to its minimal movement. Standard deviations are denoted in brackets.

| **SARS-CoV Apo Dimer** | | | **MERS-CoV Apo Dimer** | | |
|---|---|---|---|---|---|
| **Cluster** | **Upper Loop Mean $Pro_9$-$Leu_{50}$ Distance (Å)** | **State** | **Cluster** | **Upper Loop Mean $Pro_9$-$Leu_{50}$ Distance (Å)** | **State** |
| 1 | 40.8 (1.7) | Open | 1 | 37.9 (1.8) | Closed |
| 2 | 39.4 (0.9) | Open | 2 | 40.4 (1.3) | Open |
| 3 | 37.5 (0.7) | Closed | 3 | 36.0 (1.1) | Closed |
| 4 | 39.0 (1.1) | Open | 4 | 41.4 (1.1) | Open |
| 5 | 41.2 (2.2) | Open | 5 | 42.6 (1.1) | Open |
| 6 | 37.6 (1.3) | Closed | 6 | 38.5 (1.1) | Open |
| 7 | 43.6 (1.3) | Open | 7 | 39.6 (1.3) | Open |
| 8 | 41.3 (2.5) | Open | 8 | 38.7 (1.8) | Open |
| | | | 9 | 40.3 (1.2) | Open |
| | | | 10 | 40.0 (1.5) | Open |

| **Cluster** | **Lower Loop Mean $Pro_9$-$Arg_{188}$ Distance (Å)** | **State** | **Cluster** | **Lower Loop Mean $Pro_9$-$Arg_{188}$ Distance (Å)** | **State** |
|---|---|---|---|---|---|
| 1 | 33.8 (0.8) | Intermediate | 1 | 31.9 (0.4) | Closed |
| 2 | 32.7 (0.4) | Intermediate | 2 | 31.6 (0.5) | Closed |
| 3 | 33.9 (0.4) | Intermediate | 3 | 30.8 (0.6) | Closed |
| 4 | 31.9 (0.6) | Closed | 4 | 30.9 (1.8) | Closed |
| 5 | 32.6 (0.7) | Intermediate | 5 | 35.8 (2.6) | Open |
| 6 | 32.5 (0.7) | Intermediate | 6 | 34.6 (1.1) | Open |
| 7 | 32.1 (0.7) | Closed | 7 | 35.0 (2.4) | Open |
| | | | 8 | 36.3 (1.2) | Open |
| | | | 9 | 37.7 (2.3) | Open |
| | | | 10 | 35.5 (2.3) | Open |

Table S9: **Average distance from the upper or lower loop tip ($Leu_{50}$ or $Arg_{188}$) to $Pro_9$ in all UMAP and HDBSCAN clusters for the SARS-CoV-2 apo monomer.** These values were used in combination with visual inspection to categorize UMAP and HDBSCAN clusters. $Pro_9$ was chosen as a reference point due to its minimal movement. Standard deviations are denoted in brackets.

| SARS-CoV-2 Apo Monomer | | |
|---|---|---|
| **Cluster** | **Upper Loop Mean $Pro_9$-$Leu_{50}$ Distance (Å)** | **State** |
| 1 | 42.1 (1.1) | Open |
| 2 | 42.0 (1.9) | Open |
| 3 | 41.7 (2.5) | Open |
| 4 | 42.5 (1.5) | Open |
| 5 | 39.8 (1.4) | Open |
| 6 | 40.4 (1.8) | Open |
| 7 | 45.3 (1.3) | Open |
| 8 | 41.4 (2.0) | Open |
| 9 | 39.2 (2.4) | Open |
| 10 | 42.5 (4.1) | Open |

| **Cluster** | **Lower Loop Mean $Pro_9$-$Arg_{188}$ Distance (Å)** | **State** |
|---|---|---|
| 1 | 34.0 (2.6) | Intermediate |
| 2 | 39.4 (1.0) | Open |
| 3 | 35.2 (0.6) | Open |
| 4 | 36.4 (0.9) | Open |
| 5 | 37.3 (2.2) | Open |
| 6 | 36.2 (2.7) | Open |
| 7 | 35.4 (1.3) | Open |
| 8 | 33.7 (0.8) | Intermediate |
| 9 | 36.7 (1.7) | Open |
| 10 | 37.6 (1.7) | Open |
| 11 | 34.5 (1.6) | Intermediate |

### 4 Supporting Videos

- **Supporting Video S1.** An example MD trajectory from the SARS-CoV-2 $M^{pro}$ apo dimer system, rendered using Blender with the Molecular Nodes[S18] add-on. One protomer is shown in blue and the other in orange. $t = 0.5$ to 1.5 µs is shown from one of five replicas. 250 frames were used with 4 ns timesteps between each frame. One second of the video corresponds to 16 ns of simulation. Interpolation between frames was used for smoothing. The upper and lower loops can be seen changing loop states several times.

- **Supporting Video S2.** A close-up view of the active site from a SARS-CoV-2 $M^{pro}$ apo dimer trajectory, rendered using Blender with the Molecular Nodes[S18] add-on. The side chains of the upper loop (residues 44-54), lower loop (184-194), catalytic loop (162-175), and the catalytic dyad ($His_{41}$ and $Cys_{145}$) are shown in ball and stick representations. $t = 0.5$ to 0.9 µs is shown from one of five replicas. 100 frames were used with 4 ns timesteps between each frame. One second of the video corresponds to 16 ns of simulation. Interpolation between frames was used for smoothing.

- **Supporting Video S3.** Example MD trajectories from the SARS-CoV and MERS-CoV $M^{pro}$ apo dimer systems, rendered using Blender with the Molecular Nodes[S18] add-on. Protomers of each system are coloured separately. $t = 0.5$ to 0.9 µs is shown from one of five replicas for each system. 100 frames were used with 4 ns timesteps between each frame. One second of the video corresponds to 16 ns of simulation. Interpolation between frames was used for smoothing.

10.1021/acs.jcim.0c00575.

(S2) Jin, Z.; Du, X.; Xu, Y.; Deng, Y.; Liu, M.; Zhao, Y.; Zhang, B.; Li, X.; Zhang, L.; Peng, C.; Duan, Y.; Yu, J.; Wang, L.; Yang, K.; Liu, F.; Jiang, R.; Yang, X.; You, T.; Liu, X.; Yang, X.; Bai, F.; Liu, H.; Liu, X.; Guddat, L. W.; Xu, W.; Xiao, G.; Qin, C.; Shi, Z.; Jiang, H.; Rao, Z.; Yang, H. Structure of $M^{pro}$ from SARS-CoV-2 and discovery of its inhibitors. *Nature* **2020**, *582*, 289–293, DOI: 10.1038/s41586-020-2223-y.

(S3) Rut, W.; Groborz, K.; Zhang, L.; Sun, X.; Zmudzinski, M.; Pawlik, B.; Wang, X.; Jochmans, D.; Neyts, J.; Młynarski, W.; Hilgenfeld, R.; Drag, M. SARS-CoV-2 Mpro inhibitors and activity-based probes for patient-sample imaging. *Nature Chemical Biology* **2021**, *17*, 222–228, DOI: 10.1038/s41589-020-00689-z.

(S4) Ullrich, S.; Nitsche, C. The SARS-CoV-2 main protease as drug target. *Bioorganic &; Medicinal Chemistry Letters* **2020**, *30*, 127377, DOI: 10.1016/j.bmcl.2020.127377.

(S5) Berendsen, H. J. C.; Postma, J. P. M.; van Gunsteren, W. F.; DiNola, A.; Haak, J. R. Molecular dynamics with coupling to an external bath. *Journal of Chemical Physics* **1984**, *81*, 3684–3690, DOI: 10.1063/1.448118.

(S6) Gowers, R.; Linke, M.; Barnoud, J.; Reddy, T.; Melo, M.; Seyler, S.; Domański, J.; Dotson, D.; Buchoux, S.; Kenney, I.; Beckstein, O. MDAnalysis: A Python Package for the Rapid Analysis of Molecular Dynamics Simulations. Proceedings of the 15th Python in Science Conference. 2016; DOI: 10.25080/majora-629e541a-00e.

(S7) Michaud-Agrawal, N.; Denning, E. J.; Woolf, T. B.; Beckstein, O. MDAnalysis: A toolkit for the analysis of molecular dynamics simulations. *Journal of Computational Chemistry* **2011**, *32*, 2319–2327, DOI: 10.1002/jcc.21787.

(S8) Humphrey, W.; Dalke, A.; Schulten, K. VMD: Visual molecular dynamics. *Journal of Molecular Graphics* **1996**, *14*, 33–38, DOI: 10.1016/0263-7855(96)00018-5.

(S9) Meng, E. C.; Goddard, T. D.; Pettersen, E. F.; Couch, G. S.; Pearson, Z. J.; Morris, J. H.; Ferrin, T. E. ¡scp¿UCSF ChimeraX¡/scp¿: Tools for structure building and analysis. *Protein Science* **2023**, *32*, e4792, DOI: 10.1002/pro.4792.

(S10) Vögele, M.; Thomson, N. J.; Truong, S. T.; McAvity, J.; Zachariae, U.; Dror, R. O. Systematic Analysis of Biomolecular Conformational Ensembles with PENSA. 2022; https://arxiv.org/abs/2212.02714.

(S11) Virtanen, P.; Gommers, R.; Oliphant, T. E.; Haberland, M.; Reddy, T.; Cournapeau, D.; Burovski, E.; Peterson, P.; Weckesser, W.; Bright, J.; van der Walt, S. J.; Brett, M.; Wilson, J.; Millman, K. J.; Mayorov, N.; Nelson, A. R. J.; Jones, E.; Kern, R.; Larson, E.; Carey, C. J.; Polat, ; Feng, Y.; Moore, E. W.; VanderPlas, J.; Laxalde, D.; Perktold, J.; Cimrman, R.; Henriksen, I.; Quintero, E. A.; Harris, C. R.; Archibald, A. M.; Ribeiro, A. H.; Pedregosa, F.; van Mulbregt, P.; Vijaykumar, A.; Bardelli, A. P.; Rothberg, A.; Hilboll, A.; Kloeckner, A.; Scopatz, A.; Lee, A.; Rokem, A.; Woods, C. N.; Fulton, C.; Masson, C.; Häggström, C.; Fitzgerald, C.; Nicholson, D. A.; Hagen, D. R.; Pasechnik, D. V.; Olivetti, E.; Martin, E.; Wieser, E.; Silva, F.; Lenders, F.; Wilhelm, F.; Young, G.; Price, G. A.; Ingold, G.-L.; Allen, G. E.; Lee, G. R.; Audren, H.; Probst, I.; Dietrich, J. P.; Silterra, J.; Webber, J. T.; Slavič, J.; Nothman, J.; Buchner, J.; Kulick, J.; Schönberger, J. L.; de Miranda Cardoso, J. V.; Reimer, J.; Harrington, J.; Rodríguez, J. L. C.; Nunez-Iglesias, J.; Kuczynski, J.; Tritz, K.; Thoma, M.; Newville, M.; Kümmerer, M.; Bolingbroke, M.; Tartre, M.; Pak, M.; Smith, N. J.; Nowaczyk, N.; Shebanov, N.; Pavlyk, O.; Brodtkorb, P. A.; Lee, P.; McGibbon, R. T.; Feldbauer, R.; Lewis, S.; Tygier, S.; Sievert, S.; Vigna, S.; Peterson, S.; More, S.; Pudlik, T.; Oshima, T.; Pingel, T. J.; Robitaille, T. P.; Spura, T.; Jones, T. R.; Cera, T.; Leslie, T.; Zito, T.; Krauss, T.; Upadhyay, U.; Halchenko, Y. O.; Vázquez-Baeza, Y. SciPy 1.0: fundamental algorithms for scientific computing in Python. *Nature Methods* **2020**, *17*, 261–272, DOI:

10.1038/s41592-019-0686-2.

(S12) Pedregosa, F.; Varoquaux, G.; Gramfort, A.; Michel, V.; Thirion, B.; Grisel, O.; Blondel, M.; Prettenhofer, P.; Weiss, R.; Dubourg, V.; Vanderplas, J.; Passos, A.; Cournapeau, D.; Brucher, M.; Perrot, M.; Duchesnay, E. Scikit-learn: Machine Learning in Python. *Journal of Machine Learning Research* **2011**, *12*, 2825–2830.

(S13) McInnes, L.; Healy, J.; Saul, N.; Großberger, L. UMAP: Uniform Manifold Approximation and Projection. *Journal of Open Source Software* **2018**, *3*, 861, DOI: 10.21105/joss.00861.

(S14) McInnes, L.; Healy, J.; Astels, S. hdbscan: Hierarchical density based clustering. *The Journal of Open Source Software* **2017**, *2*, 205, DOI: 10.21105/joss.00205.

(S15) Klyshko, E.; Kim, J. S.-H.; McGough, L.; Valeeva, V.; Lee, E.; Ranganathan, R.; Rauscher, S. Functional protein dynamics in a crystal. *Nature Communications* **2024**, *15*, 3244, DOI: 10.1038/s41467-024-47473-4.

(S16) Sievers, F.; Wilm, A.; Dineen, D.; Gibson, T. J.; Karplus, K.; Li, W.; Lopez, R.; McWilliam, H.; Remmert, M.; Söding, J.; Thompson, J. D.; Higgins, D. G. Fast, scalable generation of high-quality protein multiple sequence alignments using Clustal Omega. *Molecular Systems Biology* **2011**, *7*, 539, DOI: 10.1038/msb.2011.75.

(S17) Shimoyama, Y. pyMSAviz: MSA visualization python package for sequence analysis. https://github.com/moshi4/pyMSAviz, 2022; https://github.com/moshi4/pyMSAviz, Version 1.2.0.

(S18) Johnston, B.; Elferich, J.; Davidson, R. B.; Zhuang, Y.; Yao, Y.; Tubiana, T.; Kunzmann, P.; Laprevote, O.; Rich; TheJeran; Autin, L.; Zwiggelaar, J.; Marson, D.; Spauszus, K. N.; Hooker, J.; Nash, J. A.; Kim, J.; Colson, L.; Copeland, H. W. BradyAJohnston/MolecularNodes: v4.1.4 for Blender 4.1. 2024; https://zenodo.

org/doi/10.5281/zenodo.6540846, Creative Commons Attribution 4.0 International.
